# Contributions of genetic variation in astrocytes to cell and molecular mechanisms of risk and resilience to late onset Alzheimer’s disease

**DOI:** 10.1101/2024.07.31.605928

**Authors:** Hyo Lee, Richard V. Pearse, Alexandra M. Lish, Cheryl Pan, Zachary M. Augur, Gizem Terzioglu, Pallavi Gaur, Meichen Liao, Masashi Fujita, Earvin S. Tio, Duc M. Duong, Daniel Felsky, Nicholas T. Seyfried, Vilas Menon, David A. Bennett, Philip L. De Jager, Tracy L. Young-Pearse

**Affiliations:** Ann Romney Center for Neurologic Diseases, Department of Neurology, Brigham and Women’s Hospital and Harvard Medical School, Boston, MA, USA; Center for Translational and Computational Neuroimmunology, Department of Neurology, and the Taub Institute for the Study of Alzheimer’s Disease and the Aging Brain, Columbia University Irving Medical Center, New York, NY, USA; Department of Psychiatry and Institute of Medical Science, University of Toronto, Toronto, Ontario, Canada; Department of Biochemistry, Emory University School of Medicine, Atlanta, Georgia, USA; Krembil Centre for Neuroinformatics, Centre for Addiction and Mental Health, Toronto, Ontario, Canada; Department of Neurology, Emory University School of Medicine, Atlanta, Georgia, USA; Rush Alzheimer’s Disease Center, Rush University Medical Center, Chicago, Illinois, USA; Harvard Stem Cell Institute, Harvard University, Cambridge, Massachusetts, USA

## Abstract

Reactive astrocytes are associated with Alzheimer’s disease (AD), and several AD genetic risk variants are associated with genes highly expressed in astrocytes. However, the contribution of genetic risk within astrocytes to cellular processes relevant to the pathogenesis of AD remains ill-defined. Here we present a resource for studying AD genetic risk in astrocytes using a large collection of induced pluripotent stem cell (iPSC) lines from deeply phenotyped individuals with a range of neuropathological and cognitive outcomes. IPSC lines from forty-four individuals were differentiated into astrocytes followed by unbiased molecular profiling using RNA sequencing and tandem mass tag-mass spectrometry. We demonstrate the utility of this resource in examining gene- and pathway-level associations with clinical and neuropathological traits, as well as in analyzing genetic risk and resilience factors through parallel analyses of iPSC-astrocytes and brain tissue from the same individuals. Our analyses reveal that genes and pathways altered in iPSC-derived astrocytes from AD individuals are concordantly dysregulated in AD brain tissue. This includes increased prefoldin proteins, extracellular matrix factors, COPI-mediated trafficking components and reduced proteins involved in cellular respiration and fatty acid oxidation. Additionally, iPSC-derived astrocytes from individuals resilient to high AD neuropathology show elevated basal levels of interferon response proteins and increased secretion of interferon gamma. Correspondingly, higher polygenic risk scores for AD are associated with lower levels of interferon response proteins. This study establishes an experimental system that integrates genetic information with a heterogeneous set of iPSCs to identify genetic contributions to molecular pathways affecting AD risk and resilience.

## 1. Introduction

Named for their star shaped morphology, astrocytes perform important functions in the brain such as delivering nutrients to neurons, establishing and maintaining synapses, clearing toxic factors such as Aβ, contributing to blood-brain barrier integrity, and promoting myelination in the white matter (reviewed in (Han et al., 2021; Linnerbauer et al., 2020)). In neurodegenerative disease, astrocytes enter an “activated” state, responding to neuronal injury with changes in gene expression that can aid in the protection and repair of damaged neurons. However, the activation of astrocytes is complex and diverse, making it challenging to characterize and define a single “reactive astrocyte” signature (reviewed in (Liddelow and Barres, 2017)).

Advancements in gene expression profiling techniques such as RNA sequencing and quantitative proteomic profiling analyses enable a more precise characterization of the molecular changes occurring in astrocytes due to environmental and genetic influences. One of the pioneering studies of astrocyte states employed a series of induced injuries for the purpose of categorizing response signatures in mice. This study identified an “A1” signature, denoting a neurotoxic phenotype triggered by specific cytokines released by microglia, and an “A2” signature, indicating a neuroprotective phenotype observed in an experimental model simulating ischemic stroke (Liddelow et al., 2017; Zamanian et al., 2012). Several subsequent studies have further classified the heterogeneous populations of astrocytes existing in the brain into subtypes using single-cell or single-nuclei RNA sequencing and identified clusters of astrocytes associated with disease (Chen et al., 2020; Fujita et al., 2024; Habib et al., 2020; Leng et al., 2022; Mathys et al., 2019; Mathys et al., 2023; Green et al., 2023).

Astrogliosis, the response of astrocytes to brain injuries and disease, is commonly associated with neurodegenerative diseases, including Alzheimer’s disease (AD). In postmortem brain tissue from individuals with early-onset AD (EOAD) and late-onset AD (LOAD), astrocytes surround amyloid beta plaques and show alterations in proteins such as GFAP (reviewed in (Chun and Lee, 2018)). Changes in morphology, upregulation of GFAP in reactive astrocytes, and increases in cytokine levels (TNF) also were observed in several AD mouse models (Domene et al., 2016; Porchet et al., 2003; Simpson et al., 2010). Importantly, multiple meta-analyses of LOAD GWAS have identified loci near genes highly expressed in astrocytes (Bellenguez et al., 2022; Kunkle et al., 2019; Lambert et al., 2013), suggesting that molecular pathways active in astrocytes contribute to AD risk and resilience. For instance, astrocytes are a major source of APOE in the brain, and the E4 allele of APOE is the strongest genetic risk for LOAD. Numerous studies have interrogated APOE biology in astrocytes, showing that APOE ε4 astrocytes exhibit morphological alterations and impaired cholesterol transport (Lin et al., 2018; Sienski et al., 2021; Tcw et al., 2022). However, aside from the extensive literature on APOE biology, relatively little is known regarding the complex contributions of AD genetic risk factors in astrocytes.

We have generated a comprehensive dataset representing astrocyte signatures derived from several genetic backgrounds. The individuals included in this dataset represented a wide range of clinical and neuropathological outcomes, including healthy brain aging, resilience to neuropathological aggregated proteins, and diagnosed AD. Using previously established astrocyte differentiation protocols, we demonstrate that induced astrocytes (iAs) derived using a direct differentiation method from iPSCs showed consistent expression of mature human astrocyte markers across various iPSC lines. We next differentiated iPSCs from 44 individuals, sourced from the Religious Order Studies and Rush Memory and Aging Project (ROSMAP (Lagomarsino et al., 2021)). These ongoing longitudinal cohort studies include participants aged 50 to over 100 years and span a spectrum of neuropathological and clinical outcomes: healthy individuals without AD neuropathology or cognitive decline, individuals with high AD neuropathology but no cognitive decline (potentially indicating resilience), and individuals with AD neuropathology and cognitive decline (Bennett et al., 2018). Astrocytes derived from these iPSC lines were deeply profiled at the RNA and protein levels. Importantly, these data are coupled to RNA and protein profiles from brain tissue from the individuals from whom they were derived, as well as to their genome sequencing data. Through various analyses, we demonstrate the utility of this resource for studying genetic contributions to AD risk in astrocytes (**Fig. 1**). We identify dysregulated proteins and pathways in astrocytes derived from individuals with AD and define the contribution of genetic risk factors to these signatures. Notably, we identify a set of proteins upregulated in astrocytes derived from individuals resilient to high neuropathological burden involved in interferon response, and a reduction in proteins suppressed by proinflammatory cytokines. These datasets and iPSC lines are available as a resource for future studies aimed at disentangling the contribution of genetic variation to cellular processes in astrocytes affecting risk and resilience to LOAD.

**Figure 1.**
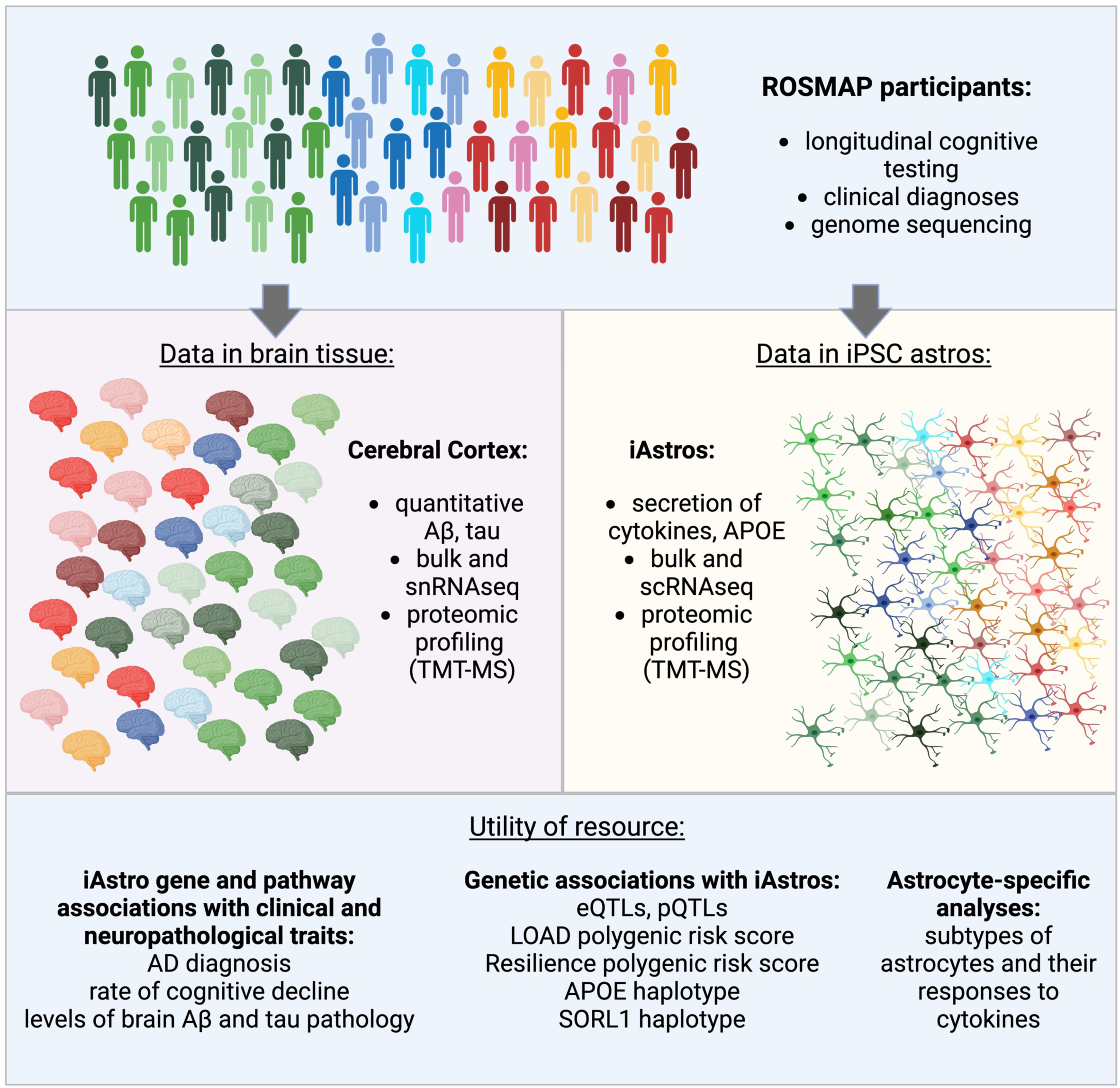
Overview of the study. Participants from the ROS and MAP cohorts that spanned the spectrum of neuropathological and cognitive outcomes were studied. In this study, IPSC-derived astrocytes derived from 44 individuals were profiled by bulk and single cell RNAseq, tandem mass-tag mass spectrometry (TMT-MS), and ELISA. These data are coupled to existing datasets derived from brain tissue of the same individuals and reported previously (Bennett et al., 2018; De Jager et al., 2018; Johnson et al., 2022; Mostafavi et al., 2018). In this study we demonstrate the utility of this resource by performing several different types of analyses to identify pathways relevant to AD in astrocytes.

## 2. Methods

### 2.1 iPSC lines

Induced Pluripotent Stem Cell lines iPSCs were generated from cryopreserved peripheral blood mononuclear cell (PBMC) samples from autopsied participants from either the Religious Order Study (ROS) or Rush Memory and Aging Project (MAP). ROS and MAP are cohort studies of aging and dementia. Each participant signed an informed consent, Anatomic Gift Act, and Repository Consent to allow their data to be repurposed, following approval by an Institutional Review Board of Rush University Medical Center. iPSCs were generated using Sendai reprogramming method as previously published (Lagomarsino et al., 2021). iPSCs undergo a rigorous quality procedure that includes a sterility check, mycoplasma testing, karyotyping, and pluripotency assays performed by the New York Stem Cell Foundation (NYSCF). iPSCs were maintained using StemFlex Medium (Thermo Fisher Scientific). All cell lines were routinely tested for mycoplasma using PCR kit (MP0035-1KT) and STR profiling to prevent potential contamination or alteration to the cell lines.

### 2.2 iPSC-derived neuron differentiation

iPSC-derived neurons (iNs) were differentiated as described in a previously published paper (Zhang et al., 2013) with minor modifications (Lagomarsino et al., 2021; Muratore et al., 2017). iPSCs were plated at a density of 95k cells/cm^2^ on plates coated with growth factor reduced Matrigel one day prior to virus transduction (Corning #354230), then transduced with three lentiviruses – pTet-O-NGN2-puro (Addgene plasmid #52047, a gift from Marius Wernig), Tet-OFUW-EGFP (Addgene plasmid #30130, a gift from Marius Wernig), and FUdeltaGW-rtTA (Addgene plasmid #19780, a gift from Konrad Hochedlinger). The cells were then replated at 200,000 cells/cm^2^ using StemFlex Medium (Thermo Fisher Scientific) and ROCK inhibitor (10μM) (D0). The media was changed to KSR media (D1), 1:1 of KSR and N2B media (D2) and N2B media (D3). On day 4, cells were dissociated using Accutase, and plated at 50,000 cells/cm^2^ using iN D4 media (NBM media + 1:50 B27 + BDNF, GDNF, CNTF (10 ng/ml, Peprotech). Doxycycline (2ug/ml, Sigma) was added from D1 to the end of the differentiation, and puromycin (5 μg/ml, Gibco) was added from D2 to the end of the differentiation. On D3, B27 supplement (1:100) (Life Technologies) was added. From D4 to the end of differentiation D21, cells were cultured iN D4 media and fed every 2-3 days.

Induced neuron protocol media:

- KSR media: Knockout DMEM, 15% KOSR, 1x MEM-NEAA, 55 μM beta mercaptoethanol, 1x GlutaMAX (Life Technologies).
- N2B media: DMEM/F12, 1x GlutaMAX (Life Technologies), 1x N2 supplement B (Stemcell Technologies), 0.3% dextrose (D-(+)-glucose, Sigma).
- NBM media: Neurobasal medium, 0.5x MEM-NEAA, 1x GlutaMAX (Life Technologies), 0.3% dextrose (D-(+)-glucose, Sigma).

### 2.3 iPSC-derived astrocyte differentiation – induced astrocytes (iAs)

iPSC-derived astrocytes (iAs) were differentiated following a previously published paper (Canals et al., 2018) with minor modifications (Lagomarsino et al., 2021). iPSCs were plated at 95k cells/cm^2^ on growth factor reduced matrigel (Corning #354230) coated plates prior to virus transduction. Then, iPSCs were transduced with three lentiviruses – Tet-O-SOX9-puro (Addgene plasmid #117269), Tet-O-NFIB-hygro (Addgene plasmid #117271), and FUdeltaGW-rtTA (Addgene plasmid #19780). The cells were then replated at 200,000 cells/cm^2^ using StemFlex Medium (Thermo Fisher Scientific) and ROCK inhibitor (10μM) (D0). The media was changed daily with Expansion Media (EM) from D1 to 3, and gradually switched from EM to FGF media from D4 to D7. On day 8, cells were dissociated using Accutase, and plated at 84,000 cells/cm^2^ using FGF media. Doxycycline (2.5μg/ml, Sigma) was added from D1 to the end of the differentiation, puromycin (1.25μg/ml, Gibco) was added on D3 of the differentiation, and hygromycin (100μg/ml, InvivoGen # ant-hg-1) was added from D4-D6 of the differentiation. From D8 to the end of differentiation D21, cells were cultured with maturation media and fed every 2-3 days.

Induced astrocyte protocol media:

- Expansion Media: DMEM/F12 (Thermo Fisher Scientific), 10% FBS, 1% N2 Supplement (Stemcell Technologies), 1% GlutaMAX (Life Technologies)
- FGF Media: Neurobasal media, 2% B27, 1% NEAA, 1% GlutaMAX, 1% FBS, 8ng/ml FGF, 5ng/ml CNTF, 10ng/ml BMP4
- Maturation Media: 1:1 DMEM/F12 and neurobasal media, 1% N2, 1% GlutaMAX, 1% Sodium Pyruvate, 5ug/ml N-1% N2, 1% GlutaMAX, 1% Sodium Pyruvate, 5μg/ml N-N54 acetyl cysteine, 5ng/ml heparin-binding EGF-like GF, 10ng/ml CNTF, 10ng/ml BMP4, 500ug/ml dbcAMP

### 2.4 iPSC-derived astrocyte differentiation – EB derived astrocytes

Embryoid body-based astrocytes (ebAs) were differentiated following a previously published paper (Tcw et al., 2017) with minor modifications. iPSCs were cultured on growth factor reduced matrigel (Corning #354230) coated 6-well plates prior to differentiation. Then, iPSCs were dissociated using collagenase and were transferred to ultra-low attachment 6 well plates with the base media (D1). The cells were then cultured with neuronal induction media (D2). The media was changed every other day by tilting the plate, allowing the embryoid bodies (EBs) to settle down (D3-7). On day 7, the EBs were transferred to a poly-ornithine/laminin (POL) coated 6 well plates using the neuronal induction media and laminin. The media was changed every other day from d7 to d14. On day 14, 1ml of rosette selection agent was added to each well in 6 well plate and incubated in 37°C incubator for 60 minutes. Next, the rosettes were removed from the plate, spun down at 300g for 3min, then plated onto a new POL coated 6 well plate using NPC media. After d21, cells were fed with human astrocyte media for 30 more days and harvested at d51.

EB astrocyte protocol media:

- Base Media: DMEM/F12 (Thermo Fisher Scientific), 1% N2 Supplement (Stemcell Technologies), 2% B27 (Life technologies)
- Neuronal induction Media: DMEM/F12 (Thermo Fisher Scientific), 1% N2 Supplement (Stemcell Technologies), 2% B27 (Life technologies), 100nM LDN 193189 (Stemgent), 10μM SB 4313542 (Stemcell technologies)
- NPC Media: DMEM/F12 (Thermo Fisher Scientific), 1% N2 Supplement (Stemcell Technologies), 2% B27 (Life technologies), 20μg/ml FGF2, 1ug/ml laminin
- Astrocyte differentiation Media: Basal astrocyte medium (ScienCell), Penn/Strep (ScienCell), FBS (ScienCell), Astrocyte Growth Supplement (ScienCell)

### 2.5 iPSC-derived microglia-like cells differentiation

iPSC-derived microglia-like cells (iMGLs) were differentiated following a previously published paper (Abud et al., 2017; McQuade et al., 2018) with minor modifications (Chou et al., 2023). iPSCs were plated on growth factor reduced matrigel (Corning #354230) using StemFlex Medium (Thermo Fisher Scientific) and ROCK inhibitor (10μM). From D0 to D12, StemDiff Hematopoietic Kit (Stemcell Technologies) was used to generated hematopoietic precursor cells (HPCs). At D12, cells were replated at 100,000 cells per 35mm well in iMGL media supplemented with 3 cytokines (IL-34 (100ng/mL, Peprotech), TGF-β1 (50ng/mL, Militenyi Biotech), M-CSF (25ng/mL, Thermo Fisher Scientific)). From D12 to D24, iMGL media with freshly added cytokines were added to the culture every other day. On D24, cells were replated at 100,000 cells per 15.6mm well with 1:1 mixture of old media and fresh iMGL media with 3 cytokines. From D24 to D37, iMGL media with freshly added 3 cytokines were added to the culture every other day. On D37, cells are resuspended in iMGL media with five cytokines, supplemented every other day until D40.

iPSC-derived microglia-like cells protocol media:

- iMGL media: DMEM/F12, 2X insulin-transferrin-selenite, 2X B27, 0.5X N2, 1X GlutaMAX, 1X non-essential amino acids, 400μM monothioglycerol, 5 μg/mL insulin
- 3 cytokines: 100 ng/mL IL-34 (Peprotech), 50 ng/mL TGF-β1 (Millitenyi Biotech), and 25ng/mL M-CSF (ThermoFisher Scientific)
- 5 cytokines: 100 ng/mL IL-34 (Peprotech), 50 ng/mL TGF-β1 (Millitenyi Biotech), and 25ng/mL M-CSF (ThermoFisher Scientific), 100ng/mL CD200 (Novoprotein), and 100ng/mL CX3CL1 (Peprotech)

### 2.6 ELISAs

48hr conditioned media was collected from d19 to d21 before harvest. 10 Proinflammatory cytokine levels (IFN-g, IL-1β, IL-2, IL-4, IL-6, IL-8, IL-10, IL-12p70, IL-13, and TNF) were measured using V-PLEX Proinflammatory Panel 1 Human kit (cat# K15049D-1).

### 2.7 Astrocyte stimulation

24hrs before harvest (d50 ebAs and d20 iAs), cells were treated with with 5ng/ml TNF and 1ng/ml IL-1β. Both TNF and IL-1β were resuspended in 0.1% BSA in PBS. After harvest, RNAseq analysis was performed to identify genes that are differentially expressed following stimulation.

### 2.8 Immunocytochemistry

At ebA d51 and iA d21, cells were washed once with PBS, then was fixed in 4% paraformaldehyde solution for 15 minutes at RT. After completing the fixation, cells were incubated in blocking buffer (2% donkey serum with 0.1% Triton X-100) for 1 hour rocking at room temperature followed by overnight incubation in primary antibody at 4°C. Then, cells are washed with PBS three times, incubated with secondary antibodies, and then washed with PBS three times. DAPI (1:1000) staining was performed during the second wash with PBS. Then, the cells were imaged using LSM710 confocal microscopy.

Antibodies for Immunocytochemistry

| Antigen host | ICC Dilution | Vendor | Catalog # |
| --- | --- | --- | --- |
| APOE goat | 1:500 | Sigma | 178479 |
| GFAP chicken | 1:1000 | Abcam | ab4672 |
| VIMENTIN mouse | 1:500 | Milipore | CBL202 |
| S100B | 1:200 | Abcam | ab52642 |
| IGFBP4 | 1:25 | Proteintech | 18500-1-AP |
| MGP | 1:200 | Proteintech | 10734-1-AP |

### 2.9 LOAD PRS calculation

AD polygenic risk scores (PRS) were calculated for 2,067 ROSMAP participants using: a) summary statistics from a genome-wide association study (GWAS) on 21,982 AD cases and 41,944 cognitively normal controls (Kunkle et al., 2019) and b) TOPMed imputed genotype data from the 530 ROSMAP participants. Polygenic risk scoring was accomplished with continuous shrinkage (PRS-cs-auto: https://github.com/getian107/PRScs), a method that uses a Bayesian approach to define effect weights rather than discrete SNP selection. The global shrinkage parameter was learnt from the data, as the source GWAS was sufficiently powered. A second set of risk scores were also calculated excluding the *APOE* region (Chr19:45,016,911-46,418,605; GRCh37). A 100 kb buffer region was applied to either end of the region as defined in the source GWAS. Any variants on chromosome 19 located in this region were removed from the summary statistics prior to processing. New Bayesian posteriors were then estimated for the remaining variants. The distribution of PRS across classes was visualized as boxplots.

### 2.10 Gene Set Enrichment Analyses (GSEA)

GSEA were performed using the desktop GSEA application (https://www.gsea-msigdb.org/gsea/index.jsp) (Subramanian et al., 2005). For associations with continuous variables such as PRS, Pearson correlation coefficients were calculated, and a signed rank file created using the −log10(p-value)*abs.value(r). For comparisons between two categories (NCI vs AD, SORL1 risk versus protective haplotype, etc), t-tests were performed for each RNA or protein, and a signed rank file created using the −log10(p-value). One thousand permutations were performed. Normalized Enrichment Scores (NES) reflect the amount of enrichment of each gene set in the top (+) or bottom (−) of the gene rank files that were normalized to account for gene set sizes. A custom gene set was generated for GSEA using RNAseq data of iA treated with IL-1β + TNF or vehicle for 24 hours, using a q-value cutoff of <0.01 for inclusion.

### 2.11 Statistical analysis

All statistical tests were performed using GraphPad Prism 9 or R. All data is shown as mean +/− SEM. Comparisons between two groups were using student’s t test, and comparisons between more than two groups were analyzed using one-way ANOVA followed by Tukey’s post-hoc test or Dunnett’s T3 multiple comparison test. Spearman correlation method (rank-order based) and Pearson correlation method (linear relationship) were used for correlation analyses.

### 2.12 Bulk RNA sequencing

High quality total RNA (eRIN > 9.0) was harvested from differentiated iAs and ebAs. Libraries were generated and sequenced to a depth of ~30 million read pairs using the GeneWiz Next-Gen sequencing service. RNAseq reads were quality trimmed to remove end base called with phred33 scores below 25 and eliminate resulting reads that are shorter than 30 bases. Trimmed reads are quantified using the Kallisto (v 0.43.1) pseudoalignment algorithm with 50 bootstraps (Bray et al., 2016).

### 2.13 Genomic data analyses

TPM Normalized expression matrices and differential expression statistics were generated using the Sleuth (v 0.30) package in R (v 4.0.3). Expression differences were tested using a Wald test after controlling for differentiation batch. Significant hits were defined by q value (FDR) < .05 and a b value > .5. Gene Ontology enrichments were tested using the clusterProfiler package (v 3.18.1) (Wu et al., 2021) in R. Enrichment dot plots were generated using ggplot2 (v 3.3.5) while the gene concept network analyses were generated using the enrichplot (v 1.12.2) package. Correlations were analyzed using custom R scripts and base correlation functions.

### 2.14 Protein Digestion

Each cell pellet was individually homogenized in 300μL of urea lysis buffer (8M urea in 10mM Tris, 100mM NaH2PO4 buffer, pH=8.5), including 5μl (100x stock) HALT protease and phosphatase inhibitor cocktail (Pierce). All homogenization was performed using a Bullet Blender (Next Advance) according to manufacturer protocols. Briefly, each tissue piece was added to Urea lysis buffer in a 1.5mL Rino tube (Next Advance) harboring 750 mg stainless steel beads (0.9-2mm in diameter) and blended twice for 5 minute intervals in the cold room (4°C). Protein supernatants were transferred to 1.5mL Eppendorf tube and sonicated (Sonic Dismembrator, Fisher Scientific) 3 times for 5s with 15s intervals of rest at 30% amplitude to disrupt nucleic acids and subsequently vortexed. Protein concentration was determined by the bicinchoninic acid (BCA) method, and samples were frozen in aliquots at −80°C. Protein homogenates (50μg) treated with 1mM dithiothreitol (DTT) at 25°C for 30 minutes, followed by 5mM iodoacetamide (IAA) at 25C for 30 minutes in the dark. Protein mixture was digested overnight with 1:100 (w/w) lysyl endopeptidase (Wako) at room temperature. The samples were then diluted with 50mM NH_4_HCO_3_ to a final concentration of less than 2M urea and then and further digested overnight with 1:50 (w/w) trypsin (Promega) at 25°C. Resulting peptides were desalted with a Sep-Pak C18 column (Waters) and dried under vacuum.

### 2.15 Tandem Mass Tag (TMT) Labeling

Peptides were reconstituted in 100μl of 100mM triethyl ammonium bicarbonate (TEAB) and labeling performed as previously described using TMTPro isobaric tags (Thermofisher Scientific, A44520) (Higginbotham et al., 2020; Ping et al., 2018). Briefly, the TMT labeling reagents were equilibrated to room temperature, and anhydrous ACN (200μL) was added to each reagent channel. Each channel was gently vortexed for 5min, and then 20μL from each TMT channel was transferred to the peptide solutions and allowed to incubate for 1h at room temperature. The reaction was quenched with 5% (vol/vol) hydroxylamine (5μl) (Pierce). All 16 channels were then combined and dried by SpeedVac (LabConco) to approximately 100μL and diluted with 1mL of 0.1% (vol/vol) TFA, then acidified to a final concentration of 1% (v/v) FA and 0.1% (v/v) TFA. Peptides were desalted with a 60mg HLB plate (Waters). The eluates were then dried to completeness. High pH fractionation was performed essentially as described (3) with slight modification. Dried samples were re-suspended in high pH loading buffer (0.07% vol/vol NH_4_OH, 0.045% v/v FA, 2% v/v ACN) and loaded onto a Water’s BEH (2.1mm x 150mm with 1.7µm beads). A Thermo Vanquish UPLC system was used to carry out the fractionation. Solvent A consisted of 0.0175% (v/v) NH_4_OH, 0.01125% (v/v) FA, and 2% (v/v) ACN; solvent B consisted of 0.0175% (v/v) NH_4_OH, 0.01125% (v/v) FA, and 90% (v/v) ACN. The sample elution was performed over a 25min gradient with a flow rate of 0.6 mL/min with a gradient from 0 to 50% B. A total of 96 individual equal volume fractions were collected across the gradient and dried to completeness using a vacuum centrifugation.

### 2.16 Liquid chromatography Coupled to Tandem Mass Spectrometry (LC-MS/MS)

All samples were analyzed on the Evosep One system using an in-house packed 15cm, 75μm i.d. capillary column with 1.9μm Reprosil-Pur C18 beads (Dr. Maisch, Ammerbuch, Germany) using the pre-programmed 21min gradient (60 samples per day) essentially as described (4). Mass spectrometry was performed with a high-field asymmetric waveform ion mobility spectrometry (FAIMS) Pro equipped Orbitrap Eclipse (Thermo) in positive ion mode using data-dependent acquisition with 2 second top speed cycles. Each cycle consisted of one full MS scan followed by as many MS/MS events that could fit within the given 2 second cycle time limit. MS scans were collected at a resolution of 120,000 (410-1600m/z range, 4×10^5^ AGC, 50ms maximum ion injection time, FAIMS compensation voltage of −45). All higher energy collision-induced dissociation (HCD) MS/MS spectra were acquired at a resolution of 30,000 (0.7m/z isolation width, 35% collision energy, 1.25×10^5^ AGC target, 54ms maximum ion time, TurboTMT on). Dynamic exclusion was set to exclude previously sequenced peaks for 20 seconds within a 10-ppm isolation window.

### 2.17 Database Searching and Protein Quantification

All raw files were searched using Thermo’s Proteome Discoverer suite (version 2.4.1.15) with Sequest HT. The spectra were searched against a human Uniprot database downloaded August 2020 (86395 target sequences). Search parameters included 10ppm precursor mass window, 0.05Da product mass window, dynamic modifications methione (+15.995Da), deamidated asparagine and glutamine (+0.984Da), phosphorylated serine, threonine and tyrosine (+79.966Da), and static modifications for carbamidomethyl cysteines (+57.021Da) and N-terminal and Lysine-tagged TMT (+304.207Da). Percolator was used filter PSMs to 0.1%. Peptides were group using strict parsimony and only razor and unique peptides were used for protein level quantitation. Reporter ions were quantified from MS2 scans using an integration tolerance of 20 ppm with the most confident centroid setting. Only unique and razor (i.e., parsimonious) peptides were considered for quantification. All RAW FILES are available on PRIDE.

### 2.18 scRNAseq of iAs and integration human snRNAseq data from the same cohorts

scRNAseq on iAs: iAs from the two donors (LPNCI individuals from the ROSMAP cohort) were pooled and harvested at d21. At d20, the cells received either vehicle (0.1% BSA in PBS) or stimulation cytokines (5ng/ml TNF and 1ng/ml IL-1β) (2.7 Astrocyte stimulation protocol). In total, 9,324 cells were used in the analysis, and roughly half of the population were cells treated with vehicle and the other half treated with cytokines. We integrated the iAs with the human snRNA-Seq data using an ‘anchor’-based integration approach via the Seurat package. First, we identified the top 2,000 highly variable genes (HVGs) across both datasets using the *SelectIntegrationFeatures* function after normalizing the data with the SCTransform method. We then used the *FindIntegrationAnchors* function to identify anchors between the two datasets based on the overlap among their nearest neighbors. Finally, we integrated the datasets using the *IntegrateData* function and visualized the integration employing a 2D UMAP projection executing *DimPlot* function from Seurat package.

## 3. Results

### 3.1 Defining the experimental system: Differentiation of iPSCs to astrocyte fate

To study the genetic contribution underlying LOAD in human astrocytes across many genetic backgrounds, it is important to choose a differentiation protocol that can reliably differentiate iPSCs into astrocyte fates without introducing excessive technical variability. Here, we selected two previously published astrocyte differentiation protocols for comparison: an embryoid body-based differentiation protocol (ebAs) and a transcription factor based induced differentiation protocol (iAs). In brief, ebAs are generated by initial differentiation into neural progenitor cells (NPCs) via dual SMAD inhibition followed by subsequent differentiation into astrocytes (**Fig. 2A**). In contrast, iAs undergo direct conversion from iPSCs to astrocytes via overexpression of the transcription factors NFIB and SOX9 (**Fig. 2A**). Both protocols have been extensively characterized previously using methods such as transcriptomics comparison with human primary astrocytes and measurement of cytokine production in response to stimulation (Tcw et al., 2017; Canals et al., 2018). Here, we differentiated iPSCs to d51 ebAs and d21 iAs based on these published protocols and confirmed that both ebAs and iAs express astrocyte markers such as glutamine synthetase (GLUL), aquaporin-4 (AQP4) and glial fibrillary acidic protein (GFAP) by immunostaining (**Fig. 2B**). We find that ebAs exhibited a relatively uniform morphology while iAs displayed diverse morphologies within each well. This diversity was consistent across different wells and cell lines. These morphological features of the iAs suggest perhaps a more heterogeneous population of astrocytes.

**Figure 2.**
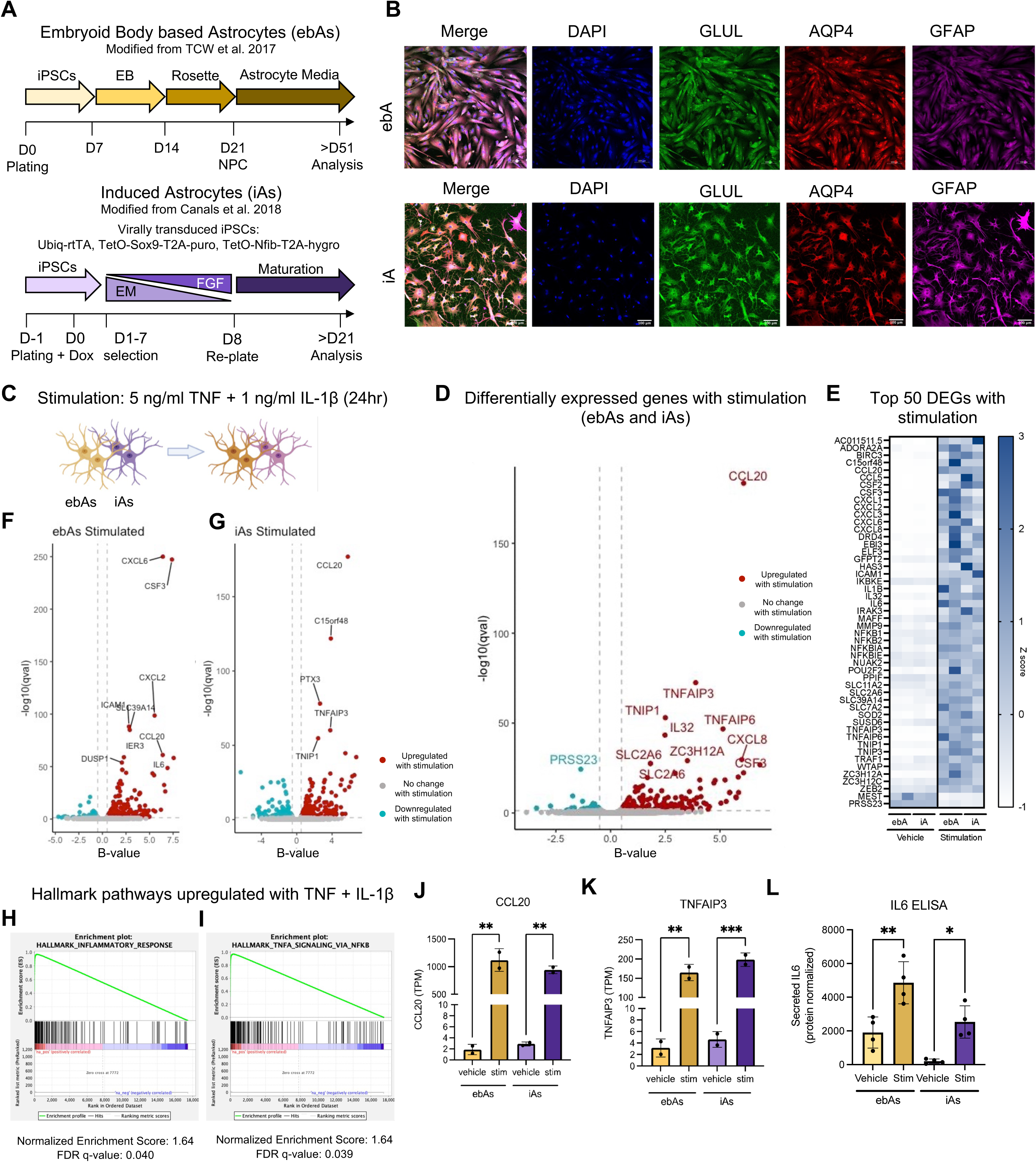
Differentiation of iPSCs to astrocyte fate using two protocols results in similar expression profile changes following stimulation with TNF and IL-1β. (A) Schematic of differentiation protocols and (B) representative immunostaining images of ebAs, iAs used in this study. Scale bars = 100 um. (C) Schematic of the stimulation paradigm. EbAs and iAs were treated with TNF (5 ng/ml) and IL-1β (1 ng/ml) for 24hours prior to lysis, and RNA samples were collected for RNAseq analysis. (D) Volcano plot of genes differentially expressed with stimulation using a combined analysis of both ebAs and iAs. DEGs reaching an adjusted p-value <0.05 are colored in turquoise (downregulated with stimulation) or red (upregulated with stimulation). (E) Heatmap of top 50 genes (z-score) that were differentially expressed with stimulation in ebAs and iAs. Separate volcano plots of genes differentially expressed with stimulation in ebAs (F) and iAs (G) also are shown. (H-I) Gene set enrichment analyses were performed following combined DEG analysis of eBAs and iAs; plots of Hallmark pathways upregulated with stimulation (FDR qval <0.05) are shown. (J-K) RNA expression levels (TPM) of genes *CCL20* (J) and *TNFAIP3* (K), which are upregulated in both ebAs and iAs with stimulation. (L) Secreted levels of IL-6 in ebA and iA cultures. Ordinary one-way ANOVA, Tukey multiple comparison test. See also Supp. Tables S1-S5).

Next, to characterize their ability to respond to cytokine stimulation, both ebAs and iAs (iPSC lines from two donors for each differentiation protocol) were stimulated with 5 ng/ml TNF and 1 ng/ml IL-1β for 24 hours followed by RNA sequencing (**Fig. 2C, Supp. Tables S1-2**). This cocktail was chosen based upon previous studies demonstrating that these factors are released by activated microglia during neuroinflammation and are capable of stimulating astrogliosis (Clausen et al., 2008; Hyvarinen et al., 2019; Rothhammer et al., 2018; Shinozaki et al., 2017; Wheeler et al., 2019). As expected, both ebAs and iAs showed numerous genes significantly up- and down-regulated in response to stimulation (adjusted q-value <0.05) (**Fig. 2D-G; Supp. Tables S3-5**). Many of the shared DEGs were upregulated genes that encode cytokines, chemokines, and genes involved in NFκB signaling. These results suggest that both ebAs and iAs can respond to cytokine stimulation, secrete cytokines, and activate immune responses in a similar manner. Gene Set Enrichment Analysis (GSEA) confirmed the generalized upregulation of Inflammation and TNFα responsive programs with stimulation (FDR qval<0.05, **Fig. 2H,I**). RNA expression levels of top DEGs such as *CCL20* and *TNFAIP3* and secreted levels of IL-6 show examples that relay that both ebAs and iAs exhibit similar effect sizes in response to stimulation (**Fig. 2J-L**).

We next characterized the gene expression profiles of ebAs and iAs compared to other iPSC-brain cell types and the extent to which iPSC-astrocytes are concordant with profiles of astrocytes present in the brain. First, we differentiated iPSC lines into neurons (iNs), astrocytes (ebAs or iAs), and microglia (iMGLs), followed by bulk transcriptomic analysis (**Fig. 3A**, **Supp. Table S6-7**). Principal component analysis (PCA) relays that ebAs and iAs have similar expression profiles to one another compared to iNs and iMGLs (**Fig. 3B**). To test the developmental maturity of cultures produced with these two protocols, we examined specific astrocyte markers identified in a previous publication (Zhang et al., 2016). That study described the transcriptomic profile of human fetal astrocytes (17-20 gestation weeks −GW) and “mature” postmitotic astrocytes in postnatal human brains (8-63 years old), thus developing astrocyte precursor cell (APC) and mature astrocyte-transcriptional profiles (Zhang et al., 2016). While ebAs and iAs both showed high expression levels of human fetal astrocyte markers compared to iNs and iMGLs, iAs also consistently expressed markers that are enriched in human “mature” astrocytes at this stage of differentiation (**Fig. 3C**). Proliferation associated markers such as *MKI67* and *TOP2A* (which are more highly expressed in immature astrocytes) and other APC-enriched markers such as *WEE1* and *VCAM1* were higher in ebAs compared to iAs (**Fig. 3D**). Both ebAs and iAs expressed canonical astrocyte markers such as *GJA1* and *SLC1A3*, but iAs expressed higher levels of human-mature astrocyte enriched markers such as *S100A1* and *WIF1* (**Fig. 3D**). If ebAs are allowed to progress beyond 51 days of differentiation, mature astrocyte marker expression is likely to increase. However, it is advantageous that the iA protocol requires a shorter differentiation time (3 weeks) compared to ebAs (>51 days: 21 days to generate NPCs, followed by an additional 30 days to generate astrocytes). An additional advantage for our purposes is that the iA protocol involves a selection process which reduces the variability that can be observed when the differentiation efficiency varies across iPSC lines. Therefore, while both protocols are useful, the iA protocol was utilized for subsequent experiments in this study.

**Figure 3.**
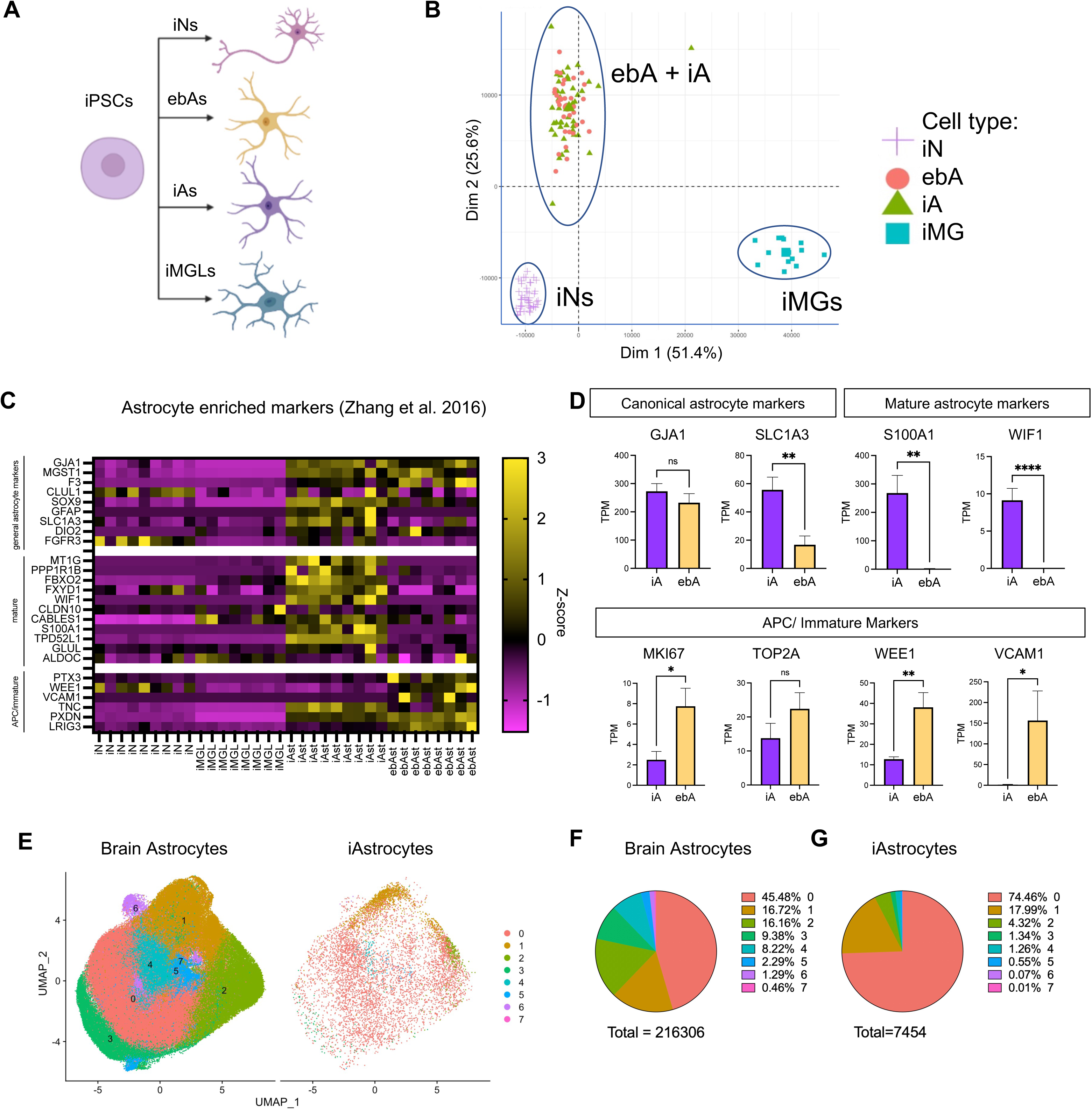
Both ebAs and iAs express astrocyte markers; iAs express higher levels of mature astrocyte markers. (A) Schematic of the experimental design. A subset of iPSCs lines were differentiated into neurons (iN), EB astrocytes (ebA), induced astrocytes (iA), and microglia (iMGL) and bulk RNAseq performed. (B) PCA plot of iNs, ebAs, iAs, AND iMGLs. (C,D) Heatmap of relative RNA levels (z-score) of canonical astrocyte markers, fetal and mature astrocyte markers and bar graphs of a subset of these markers (markers identified in (Zhang et al., 2016)). Data in (D) show mean +/− SEM, n=8 lines per group. Unpaired student’s t test (two-tailed). (L-N) Single cell RNAseq comparison between brain astrocytes and iAs. (E) UMAP plot shows seven clusters in brain astrocytes and iAs. (F,G) Pie chart showing the distribution of each cluster in (M) brain astrocytes and (N) iAs. See also Supp. Tables S6, S7.

We next aimed to interrogate whether the iA population generated is consistent with subtypes of astrocytes found in the human brain. To this end, we generated single cell RNAseq (scRNAseq) data from iAs in two genetic backgrounds. We then compared the profiles of these cells to astrocytes found in adult brain tissue. We utilized previously described single nucleus RNAseq (snRNAseq) data from human brain tissue across 424 aged participants in the ROS and MAP cohorts (average age at death, 89 years, (Fujita et al., 2024; Green et al., 2023). From this snRNAseq, we extracted data for those cells in the astrocyte cluster (216,306 cells) and harmonized these data with scRNAseq from iAs (7,454 cells). This analysis resulted in eight clusters, and each of these contained brain astrocytes and iAs to different degrees (**Fig.3E-G**). 92% of iAs were found in clusters 0 and 1, clusters which represented 62% of the brain astrocyte population (**Fig.3F,G**). Gene ontology (GO) analyses of marker genes for cluster 0 revealed leading terms of aerobic electron transport chain (GO:0019646), cytoplasmic translation (GO:0002181), and ubiquitin ligase inhibitor activity (GO:1990948) while cluster 1 was represented by regulation of response to stimulus (GO:0048583), transforming growth factor beta receptor activity (GO:0005024), and positive regulation of cell migration (GO:0030335). These data suggest that iAs share profiles aligning with subsets of astrocytes found in the adult brain.

### 3.2 Single cell RNAseq reveals three distinct astrocyte subtypes in iA cultures with overlapping but unique responses to treatment with IL-1β and TNF

Recent studies have expanded from the traditional binary division of ‘neurotoxic’ and ‘neuroprotective’ astrocyte signatures and utilized scRNAseq to identify multiple clusters within astrocytes in the human brain (Fujita et al., 2024; Green et al., 2023; Hasel et al., 2021; Leng et al., 2022; Mathys et al., 2023). Activation and release of cytokines in response to injury or other perturbations has been one of the key features of astrocytes. However, there is no simple way to define and categorize “reactive” signatures (Escartin et al., 2021), perhaps due in part to the heterogeneity in astrocyte populations. To comprehensively describe the transcriptional profile of reactive astrocytes within our model system, we first captured the signatures of stimulated astrocytes by treating iAs from nine genetic backgrounds with TNF and IL-1β for 24 hours. This was followed by bulk RNA sequencing and cytokine measurements. Astrocytes from all nine lines responded by secreting similar levels of IL-6 (**Supp. Fig. 1A-C**). Importantly, for analyses arising in subsequent sections, the generation of this RNAseq dataset allowed us to develop two gene sets that define the transcriptional responses of iAs to activation: 1) genes upregulated with stimulation (q-value<0.01) and 2) genes downregulated with stimulation (q-value<0.01) (**Supp. Tables 8,9**).

Next, we utilized the iA scRNAseq dataset described in the previous section to further explore heterogeneity in iA cultures both at baseline and in response to IL-1β + TNF treatment for 24 hours (**Fig. 4A,B**). Roughly half of the analyzed cells consisted of astrocytes treated with vehicle and the other half were those treated with cytokines (**Fig. 4C**), and a similar number of cells from each donor were represented across all clusters (**Fig. 4D**). Stimulated and unstimulated iAs separated cleanly along the UMAP1 axis (**Fig. 4C**). Examination of the top upregulated genes with stimulation (for ex., several *CCL*s, *CXCL*s, and *TNFAIP*s) revealed consistent induction of chemokines and related genes across clusters of stimulated iAs (**Fig. 4E,H,I; Supp. Table 10**). Pathway analysis of downregulated genes revealed enrichment in genes involved in regulation of locomotion (GO:0040012; FDR = 9E-7) and fatty acid oxidation (GO:0019395; FDR = 0.01), among others (**Fig. 4F**).

**Figure 4.**
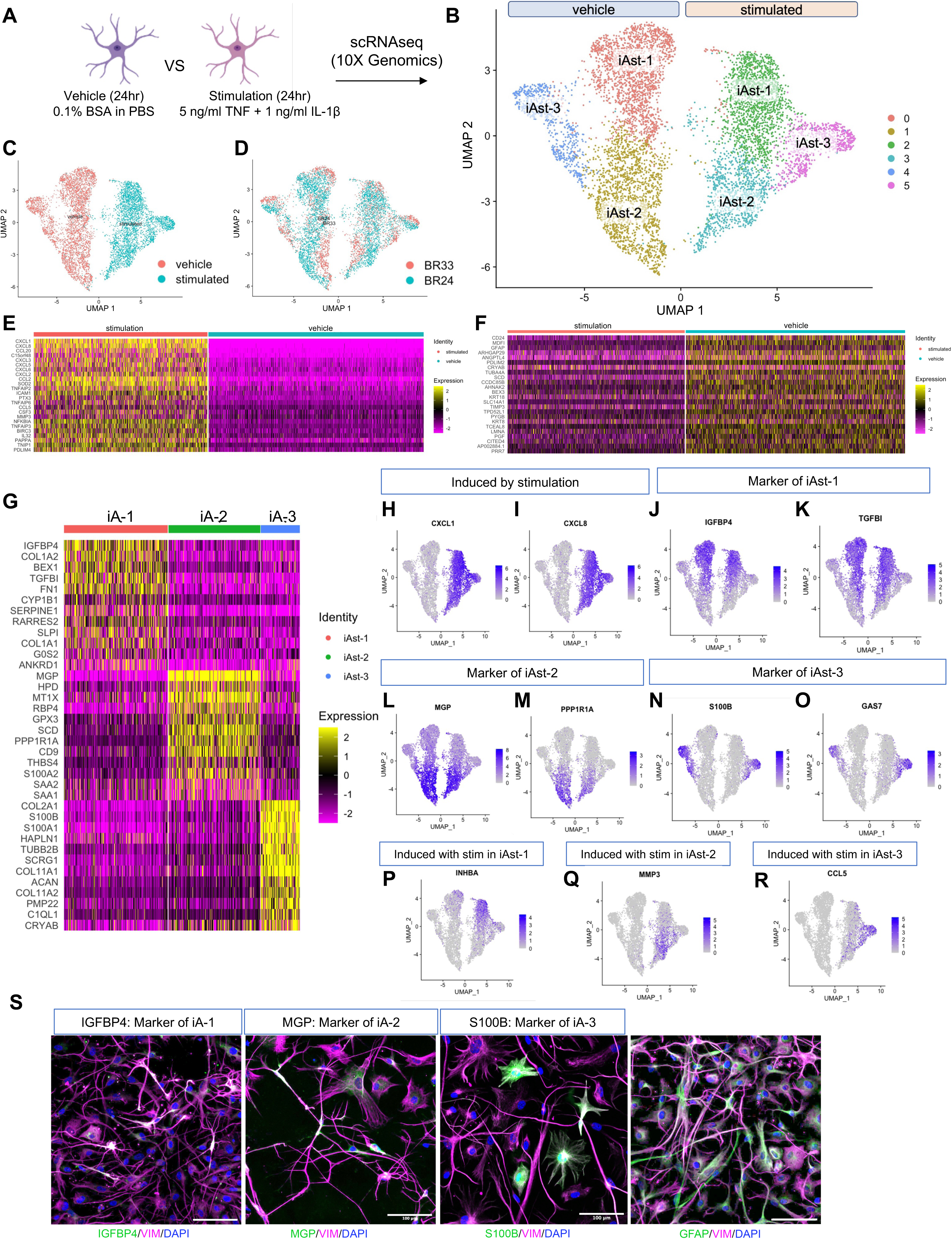
Three subtypes of astrocytes are present in iA cultures that respond to stimulation with overlapping but distinct expression changes. (A) Schematic of the experimental design. iAs from two donors were analyzed at d21 of differentiation by single cell RNAseq. For the last 24 hours of culture prior to harvest, the cells were treated with either vehicle (0.1% BSA in PBS) or cytokines (5ng/ml TNF and 1ng/ml IL-1β). (B-D) UMAP plot shows iAs cluster separately based on treatment status at the single cell level. In total, 9,324 cells were analyzed with 5,113 vehicle treated cells and 4,211 cells stimulated with cytokines (C). Cells of each genetic background (BR24, BR33) appeared in all clusters (D) Heatmap of expression levels of top genes upregulated (E) and downregulated (F) with stimulation. Six clusters were identified across vehicle and stimulation conditions: the three clusters in vehicle treated conditions (iA-1, iA-2, iA-3) were readily identified by expression profile in stimulation conditions. (G) Heat map showing markers of iA-1, -2, and -3. (H-R) UMAP plots of examples of genes marking specific clusters. (S) Immunostaining of cluster markers in unstimulated iA cultures. Scale bar = 100um. See also Supp. Tables S8, S9.

Three clusters of iAs were identified based upon global expression profiles within the vehicle-treated population, and these clusters clearly mapped to three clusters in the stimulated population (**Fig. 4B**). Differential gene expression (DEG) analyses were performed to compare gene expression between clusters (**Supp. Fig. 2A-C**). The top 12 genes driving membership in these three subpopulations are shown (**Fig. 4G**): for example, *IGFBP4* and *TGFB1* expression were strongly enriched in iAst-1, *MGP* and *PPP1R1A* expression were enriched in iAst-2 and *S100B* and *GAS7* expression were enriched in iAst3 (**Fig. 4J-O**). Interestingly, while many genes upregulated with stimulation were shared by all three iAst subtypes (**Fig. 4H,I**), some gene expression changes were specific to only one of the subtypes. For example, *INHBA* was upregulated in iAst-1, *MMP3* was upregulated in iAst-2 and *CCL5* was upregulated in iAst-3 (**Fig. 4P-R**). **Supp. Table 11** lists the markers identified for each cluster from this analysis.

Immunostaining for markers of each subtype revealed morphological differences between cells expressing IGFBP4 (iAst1), MGP (iAst2) and S100B (iAst3) (**Fig. 4S**). The iAst-1 subcluster was enriched for pathways including “ubiquitin protein ligase binding”, “protein folding”, “immune response”, “mitochondrion organization” and “aerobic respiration”. The iAst-2 subcluster showed enrichment for pathways including “cell migration”, “amyloid-beta binding” and “extracellular matrix”. The iAst-3 subcluster may reflect a level of reactive astrocyte state even in the absence of cytokine stimulation. This subcluster was enriched for the terms “MHC protein complex assembly” and “antigen processing and presentation of exogenous peptide antigen via MHC class II”.

### 3.3 Congruence of gene expression profiles between astrocyte cultures and brain

We then compared iAs derived from a large number of iPSC lines derived from the ROS and MAP cohorts to determine the capacity of this experimental system to capture the impact of genetic variation on iA biology. Approximately one third of the individuals had both a clinical and pathological diagnosis of Alzheimer’s disease (AD) while the rest were not cognitively impaired (NCI). Here, iPSC lines from 44 individuals (27 NCI and 17 AD) were differentiated to astrocyte fate using the iA protocol and the resultant iAs analyzed (**Supp. Fig. 3,4, Supp. Table 12**). To validate the reproducibility of the data, we first performed RNAseq analysis on nine lines as a discovery (pilot) dataset followed by a validation (main) cohort using all 44 lines which includes cells from the same nine lines used in the pilot cohort (**Fig. 5A**). RNAseq data from all samples, including the 9 replicates, were integrated and plotted in the same tSNE space (**Fig. 5B**). This analysis showed that the transcriptomic profiles of the nine overlapping individuals cluster together according to background genotype, which raises confidence that differences observed in expression profiles across lines are primarily intrinsically encoded within the genome and not due to technical noise.

**Figure 5.**
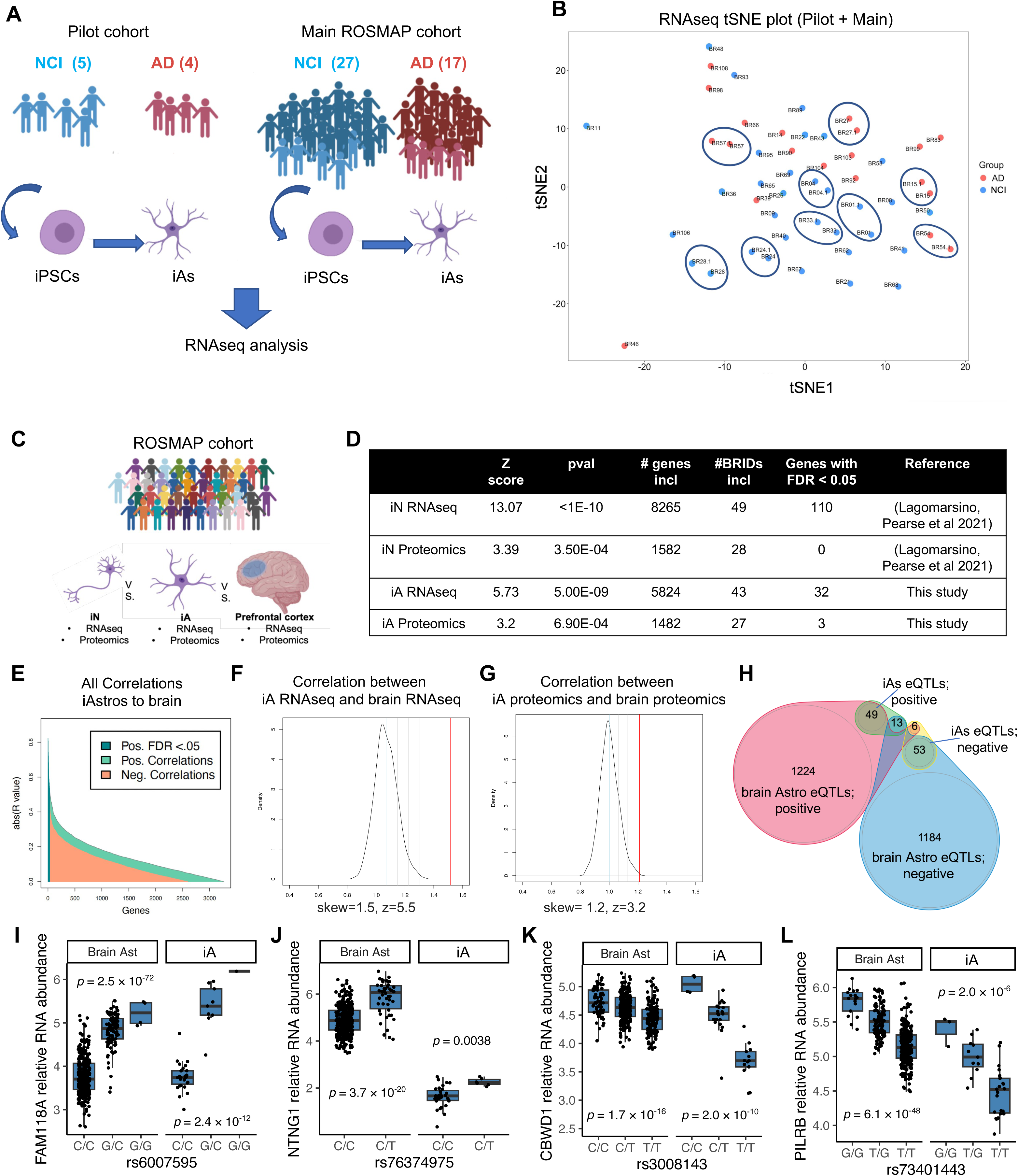
Congruence of gene expression profiles between astrocyte cultures and brain tissue from matched donors. (A) RNAseq was performed on a pilot cohort of iAs generated from 9 ROSMAP iPSC lines followed by a larger cohort of iAs generated from 44 ROSMAP iPSC lines, which also contained the original 9 iAs. (B) Following batch correction, iA RNAseq profiles cluster together based upon genetic background. (C) RNAseq and proteomics profiles from iAs and corresponding brain tissue (mPFC) from the same individuals were compared. For each gene in the upper half of percentage variance, the Pearson correlation (r value) was calculated between iAs and brain across human subjects. (D) Table of summary statistics determining congruence between gene expression profiles (RNAseq and Proteomics) between ROSMAP brain tissue and ROSMAP iNs (Lagomarsino et al. 2021) and ROSMAP iAs. (E) Waterfall plot of r values for 5,824 genes included in the comparison between RNAseq profiles between iAs and brain tissue. (F,G) Positive skew of all r values as a population was measured by dividing the sum of the highest 100 r values by the absolute value of the lowest 100 r values. Density plot of null distribution of skews is shown. The measured skew (vertical red line) is significantly higher than what would be expected by chance. (H) Venn diagram showing overlap in eQTLs between iAs and brain astrocytes calculated from single nucleus RNAseq (snRNAseq) data from (Fujita et al., 2024)). Directionality of associations with SNPs are relevant to congruence: “positive” eQTLs are SNPs that are associated with higher expression of the associated gene (examples shown in I,J) while “negative” eQTLs are SNPs that are associated with reduced expression (as in K,L). See also Supp. Tables S10-S12.

One of the unique strengths of using iAs from the ROSMAP cohort is the ability to run a direct test of overlap between measurements from iAs and from the postmortem brains from whom these iPSCs were derived (**Fig. 5C**). To assess the relevance of our model to the human brain we quantified the level of transcriptional correlation between RNA-seq gene expression profiles from iAs and the medial prefrontal cortex (PFC; BA9/46) of the same individuals. Previously we have performed a similar analysis comparing iPSC-derived neurons (iNs) from the ROSMAP cohort with the postmortem brain from whom our iPSCs were derived and observed a high congruence between iNs and brain RNAseq ((Lagomarsino et al., 2021), **Fig. 5D**). For this analysis, the dataset was filtered to compare only genes in the upper half of variance across iA samples to reduce the false correlation of technical noise. We calculated Pearson correlations (r values) for individual gene-level RNA detected in both iAs and brain (5,824 genes in total) across all 44 distinct genotypes. If there were no physiological congruence between the two datasets, then we expect to see a roughly equal number of positive correlations as we see negative correlations by chance. We rationalized that any genetically encoded congruence between iAs and brain would be observed as a positive skew in the distribution of all r values (more positive r values than negative), since an individual negative correlation would have suspect biological value. We calculated skew in our correlation analyses by dividing the sum of the highest 100 r values by the absolute value of the lowest 100 r values though other skew calculations produced similar results. To determine whether the population level correlation skew was greater than would be expected by chance, we permuted the samples over 1000 iterations to generate a null distribution of correlations that would be expected by chance. We indeed observed a positive skew of all brain-to-iA gene correlations that was significantly higher than what would be expected by chance with a z score of 5.5 and a p value of 1.8E-8 (**Fig. 5D-G**). These data suggest that this experimental system captures a measurable subset of the intrinsically encoded genetic differences between individuals and that there is a quantifiable congruence between iAs and brain tissue of the individuals from whom they were derived. A similar analysis using the proteomic data from both iAs and postmortem brain also showed a positive skew in correlations that was greater than would be expected by chance (z=3.2, p=6.8E-4) (**Fig. 5D, Supp. Table 13,14**).

Congruence in gene expression profiles between iAs and human brain tissue reflects the consistency of the impact of genetic variation on gene expression between both tissues. Expression QTL analyses identify single nucleotide polymorphisms (SNPs) that are associated with levels of expression of nearby (cis-eQTL) or distant (trans-eQTL) genes. Here, we aimed to examine astrocyte eQTLs identified in brain tissue to determine if they had similar associations in iAs. A recent study used single nucleus RNAseq (snRNAseq) data from human brain tissue across 479 ROSMAP participants to identify cell type-specific eQTLs (Fujita et al., 2024). In astrocytes, 2,529 eQTLs were identified. Of these, 102 eQTLs were replicated in our relatively small cohort of iAs (**Fig. 5H-L**) We anticipate that as more genetic backgrounds are added to iPSC cohorts, additional eQTLs from the brain will be validated *in vitro*. Thus, even with the obvious differences between iN and iA cultures and brain tissue, we observe a readily detectable and significant overlap of genetic regulation. Furthermore, the congruence between iNs/iAs and the brain can be attributed to both identified eQTLs and yet-to-be-identified genetic variants that regulate gene expression. Having demonstrated the striking resemblance of this cohort of astrocytes to signatures found in the human postmortem brain, the subsequent sections showcase the diverse applications of this resource. These include pinpointing pathways disrupted in AD and resilient individuals, as well as unraveling the molecular intricacies of established AD genetic risk factors.

### 3.4 Identification of pathways and proteins altered in human LOAD astrocytes

Genetic and neuropathology studies of AD suggest that astrocytes play a key role in risk and resilience to AD. Here, we aimed to interrogate whether genetically encoded risk for AD has measurable effects on astrocyte biology in our cellular system. To address this question, we first took an unbiased approach to identify genes, proteins, and/or pathways altered in LOAD astrocytes by utilizing the bulk RNA-seq and proteomic profiling data from the ROSMAP iA cultures (**Fig. 6A)**. Differential gene and protein expression (DEG, DEP) comparing iAs derived from AD iPSC lines to those derived from NCI lines revealed sets of genes that showed concordant RNA- and protein-level up- and down-regulation in AD iAs (**Fig. 6B,C, Supp Tables 15**). One particularly interesting example of a gene upregulated at both the RNA and protein level is CD38 (**Fig. 6C**), an enzyme that has been shown by multiple studies to be upregulated in reactive astrocytes and play an important role in mediating stress-induced inflammatory responses (Meyer et al., 2022; Wang et al., 2024).

**Figure 6.**
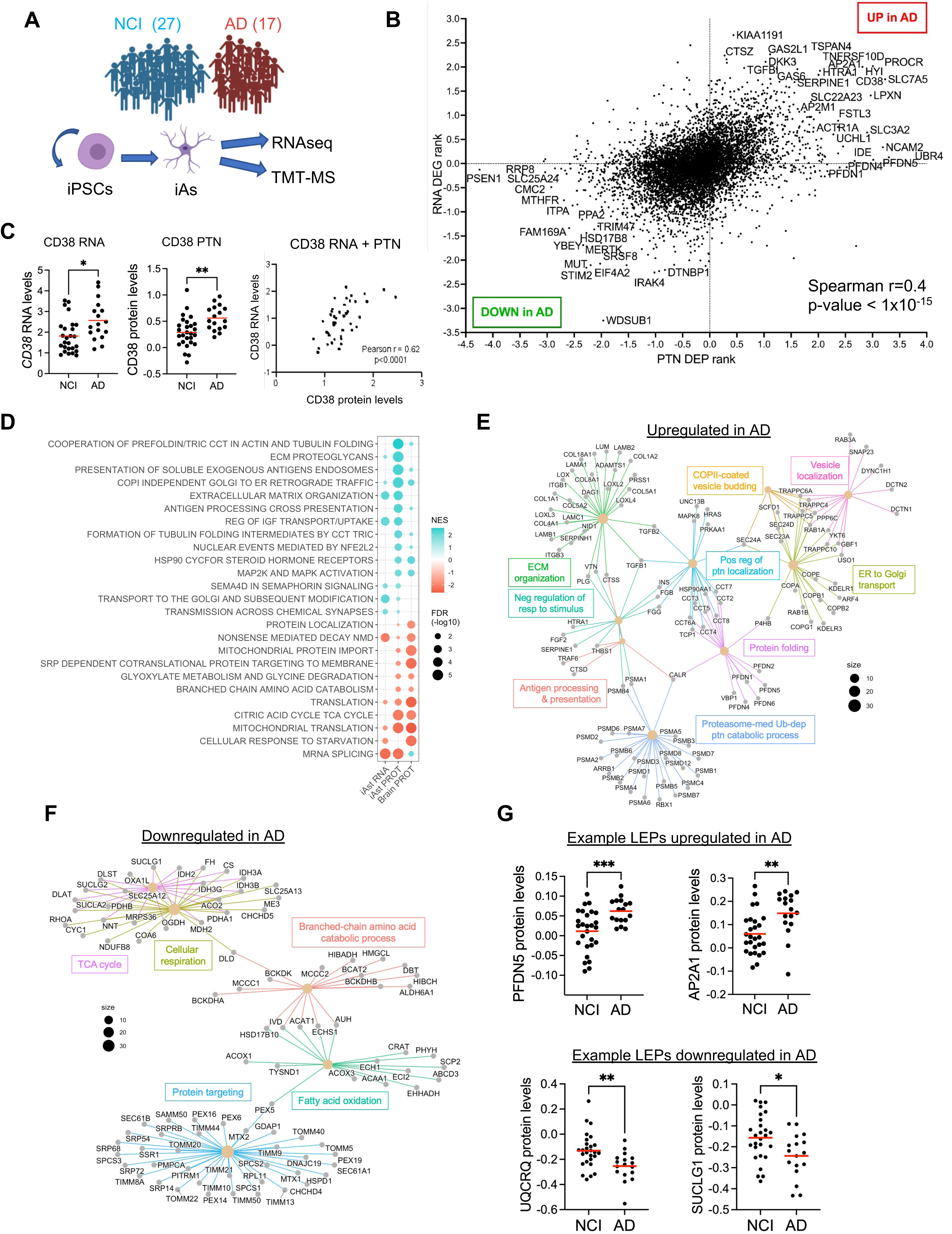
Identification of pathways and proteins altered in human LOAD astrocytes. (A) Schematic of the experimental design. iAs generated from 44 individuals were differentiated to d21. D21 iA lysates were analyzed via RNAseq and tandem mass tag mass spectrometry (TMT-MS) proteomics. B) Differential expression at the gene and protein level was analyzed and ranked by calculating the −log10(qvalue): higher DEP/DEG values being associated with higher expression in AD astrocytes compared to NCI astrocytes. Overall, a positive correlation was observed between differences (AD vs NCI) in RNA and protein levels at a gene level (Spearman correlation coefficient r = 0.4; p-value <1×10^−15^). Each dot represents data for a single gene. Some factors are AD astrocyte-enriched (upper right section) and others are NCI astrocyte-enriched (lower left section) in both the RNAseq and the proteomics data. (C) Examples of genes that are differentially expressed in both RNA and protein datasets comparing NCI and AD astrocytes. (D) Gene set enrichment analyses (GSEA) was performed from the. RNA and protein AD vs NCI analysis. Shown are pathway enrichment dot plots representing biological gene cohorts that are significantly overrepresented in either the NCI astrocytes (POS) or AD astrocytes (NEG) in RNAseq and proteomics data. In parallel, the same analysis was performed on a previously published dataset of proteomic profiles of ROSMAP brain profiles (ref), and results of concordant pathways enriched in AD or NCI brain tissue are shown. Dot color represents the number of genes passing correlation cutoff that were in the gene set while size indicates q-value. All pathways shown had a q-value < 0.02. (E,F) Leading edge proteins identified through pathway analyses in proteomics data were utilized to generate gene concept networks (GCN). Shown is the GCN of proteins upregulated in AD (E) and downregulated in AD (F); and scatter plots of levels of example leading edge proteins (LEPs) from the GSEA analysis (G). For all panels, *p<0.05, **p<0.01, ***p<0.005. See also Supp. Tables S10-16.

Pathway analyses using GSEA similarly revealed concordant pathways upregulated in AD iAs at both the RNA and protein levels (**Fig. 6D, Supp Tables 16-17**). To determine if any of these pathways also are altered in AD brain tissue, we performed the same GSEA on a previously published proteomics dataset of brain tissue from 340 ROSMAP participants (Johnson et al., 2022). Intriguingly, a subset of pathways altered in ROSMAP iAs showed concordant dysregulation in human dorsal-lateral prefrontal cortex (DL-PFC) (**Fig. 6D**, **Supp Table 18**). Gene concept networks of the leading-edge proteins driving association with pathways from proteomics data are shown to highlight the pathways and key genes that are upregulated (**Fig. 6E**) and those that are downregulated (**Fig. 6F**) in AD iAs compared to NCI iAs. Several interrelated pathways upregulated in AD iAs involve ER-Golgi trafficking and protein folding such as PFDN5 and AP2A1 (**Fig. 6D,E,G**) while downregulated pathways involve mitochondrial function and fatty acid oxidation such as UQCRQ and SUCLG1. (**Fig. 6D,F,G**).

### 3.5 Astrocytes from individuals resilient to the accumulation of AD neuropathology show elevated levels of proteins involved in interferon gamma signaling under basal conditions

In the aging human population, there are a subset of individuals who do not develop Alzheimer’s dementia despite having a high enough burden of amyloid plaques and tau tangles to warrant a neuropathological diagnosis of AD (reviewed in (Young-Pearse et al., 2023)). One potential factor that may contribute to this resilience to high plaque burden may lie in the ability of glial cells to respond efficiently to accumulating Aβ and pathogenic tau. We next aimed to utilize the established datasets to examine whether intrinsic, genetically encoded programs in astrocytes may contribute to protection in individuals with a neuropathological diagnosis of AD who were cognitively unimpaired. To this end, we compared iA protein expression profiles from individuals with a clinical and neuropathological diagnosis of AD to iAs from those who were cognitively unimpaired but who received a postmortem neuropathological diagnosis of AD (high pathology not cognitively impaired; HP-NCI). GSEA revealed that HP-NCI iAs from “resilient” individuals expressed higher levels of proteins associated with interferon response and oxidative phosphorylation (**Fig. 7A-D**, **Supp. Fig. 5, Supp. Tables 19,20**). Leading edge proteins driving this association are known target genes of interferon signaling (ISG15, ISG20, IFIT1, STAT2) as well a regulator of interferon signaling, TRIM21 (Manocha et al., 2014). Furthermore, HP-NCI iAs secreted a higher level of IFN-γ relative to LP-NCI or AD iAs (**Fig. 7E**, **Supp. Fig. 6**). These analyses raised the possibility that HP-NCI iAs may exist in a state of low-level activation. To investigate this, we applied our previous custom gene sets, which define the transcriptional responses of iAs following TNF + IL-1β stimulation (**Supp. Fig. 1, Supp. Tables 8,9**), to our HPNCI vs AD datasets. While the “upregulated with IL-1β + TNF stimulation” gene set was not enriched in HP-NCI astrocytes, genes that were downregulated with stimulation were significantly downregulated in HP-NCI iAs compared to AD iAs at both RNA and protein level (for RNAseq data FDR q-value <1E-5; FDR for proteomics data FDR q-value=0.027; **Fig. 7F**). One example is the gene DKK3, which is downregulated at both the RNA and protein levels in HPNCI iAs compared to AD iAs, as well as at the protein level in the human brain (**Fig. 7G**). These findings suggest that genetic resilience factors in HPNCI iAs may result in an iA state that is “primed” to respond more efficiently than AD iAs to environmental cues. Future experiments beyond the scope of the current study are necessary to further probe this hypothesis.

**Figure 7.**
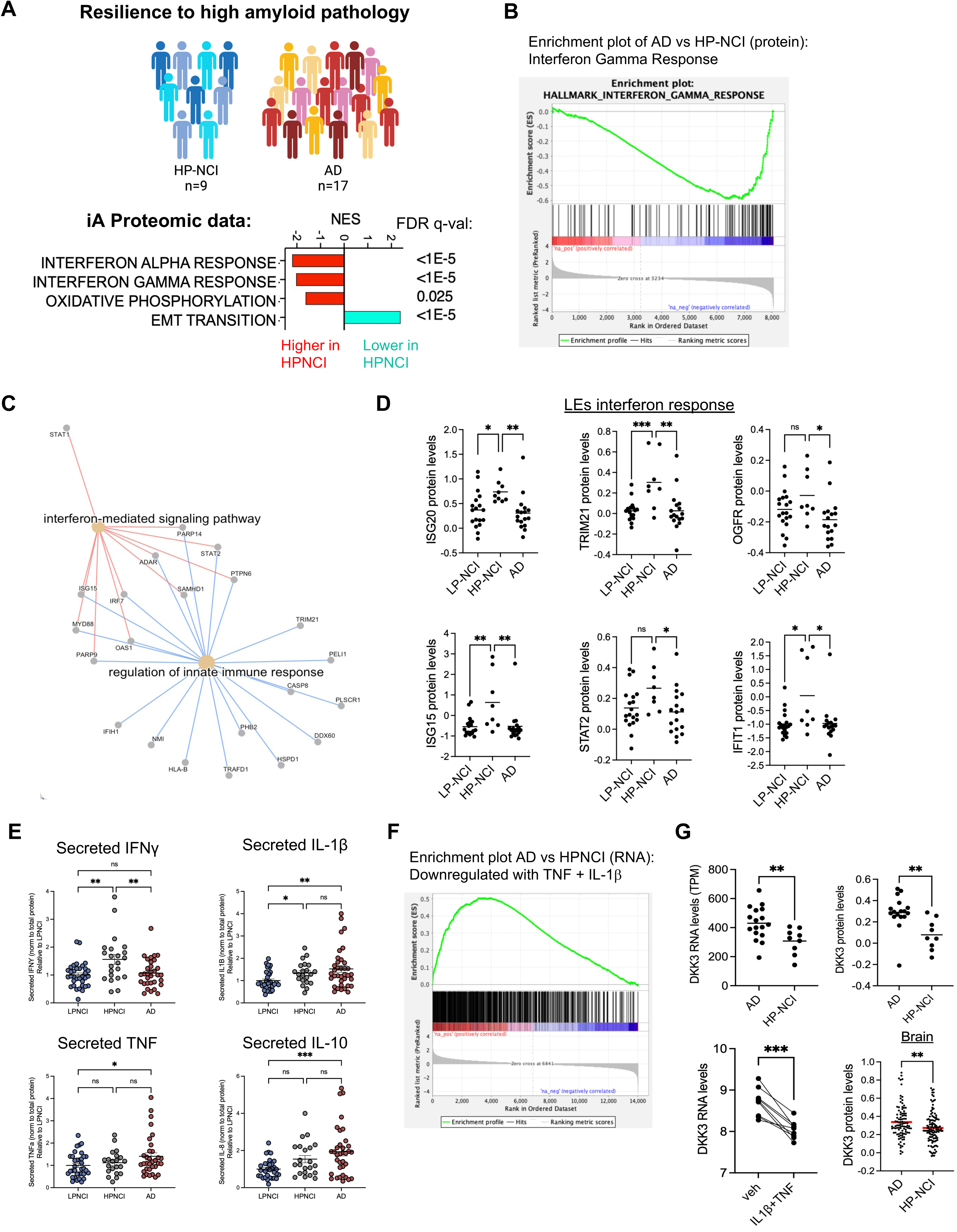
Examination of astrocytes from individuals resilient to the accumulation of AD neuropathology. (A-D) Proteomic profiles of ROSMAP iAs were examined, comparing iA from individuals with a neuropathological diagnosis of AD who did not have a clinical diagnosis of dementia (HP-NCI) to those with both a clinical and neuropathological diagnosis of AD. Differential protein analysis was performed followed by GSEA using the Hallmark gene sets (A,B). Leading edge (LE) proteins driving the association of higher interferon response in HP-NCI iAs were used to generate a gene concept network (C), and levels of example LEPs across categories are shown (D). (E) Secreted levels of IFNγ, IL-1β, TNF, and IL-10 were measured via ELISA and categorized by diagnosis. Each measurement is normalized to total protein and values shown are relative to LPNCI individuals. One-way ANOVA with Tukey’s multiple comparison’s test, n=2 wells per cell line (F) GSEA using a custom gene set of genes upregulated and downregulated in iAs with TNF + IL-1β stimulation was employed, and DEPs downregulated in HP-NCI compared to AD iAs were enriched in genes downregulated with cytokine stimulation in iAs as baseline. (G) An example leading edge gene showing downregulation in HP-NCI iAs at both the RNA and protein level, downregulation with cytokine stimulation, and downregulation in HP-NCI brain tissue at the protein level. For each graph, each dot represents data from one participant; t-tests performed for each comparison, ** p<0.01; *** p<0.0001. See also Supp. Table 17.

### 3.6 Interrogation of the contributions of genetic variants to alterations of molecular pathways in astrocytes

Genetic contributions to risk and resilience to AD within each diagnosis group are diverse, thus, an alternative, complementary approach to comparing across diagnostic categories is to interrogate the contributions of specific genetic variants to alterations of molecular pathways. Here, we performed association studies of RNA and/or protein profiles with LOAD polygenic risk score, *APOE* and *SORL1* haplotype.

In our previous study of neurons derived from this ROSMAP iPSC cohort, we found that while no single LOAD-associated allelic variant showed significant association with Aβ or tau levels, polygenic risk score for LOAD was significantly associated with the Aβ profile produced by iNs (Lagomarsino et al., 2021). Here, we examined which genes and pathways are associated with LOAD PRS within astrocytes. LOAD PRS was calculated based upon the summary statistics for LOAD (Bellenguez et al., 2022) using a Bayesian approach. In the ROSMAP iA cohort studied here, LOAD polygenic risk score was elevated in AD cases, however, there was a large amount of overlap between AD and NCI, highlighting different contributions of known genetic risk factors across individuals (**Fig. 8A**). *APOE* is the strongest genetic risk factor in AD, with the presence of the ε4 genotype conferring 3-5-fold higher risk to AD, and therefore has a strong influence on LOAD PRS (**Fig. 8A**). Interestingly, high LOAD PRS was associated with reduced transcripts and proteins in the interferon response pathway (**Fig. 8B-E, Supp Table 21-24**). However, APOE ε4 was not associated with a reduction in proteins in the interferon response pathway (**Fig. 8F**), suggesting that this association with PRS was independent of the *APOE* locus. In contrast, both high LOAD PRS and *APOE* ε4 were concordantly associated with reduced proteins relating to oxidative phosphorylation and elevated levels of proteins involved in protein secretion and hypoxia (**Fig. 8E,F, Supp Table 25,26**). A deeper analysis of the consequences of *APOE* ε4 on astrocyte biology in a large cohort of isogenic iAs that are *APOE* ε4/ ε4 vs ε3/ ε3 can be found here (Blanchard et al., 2022; Lin et al., 2018; Tcw et al., 2022).

**Figure 8.**
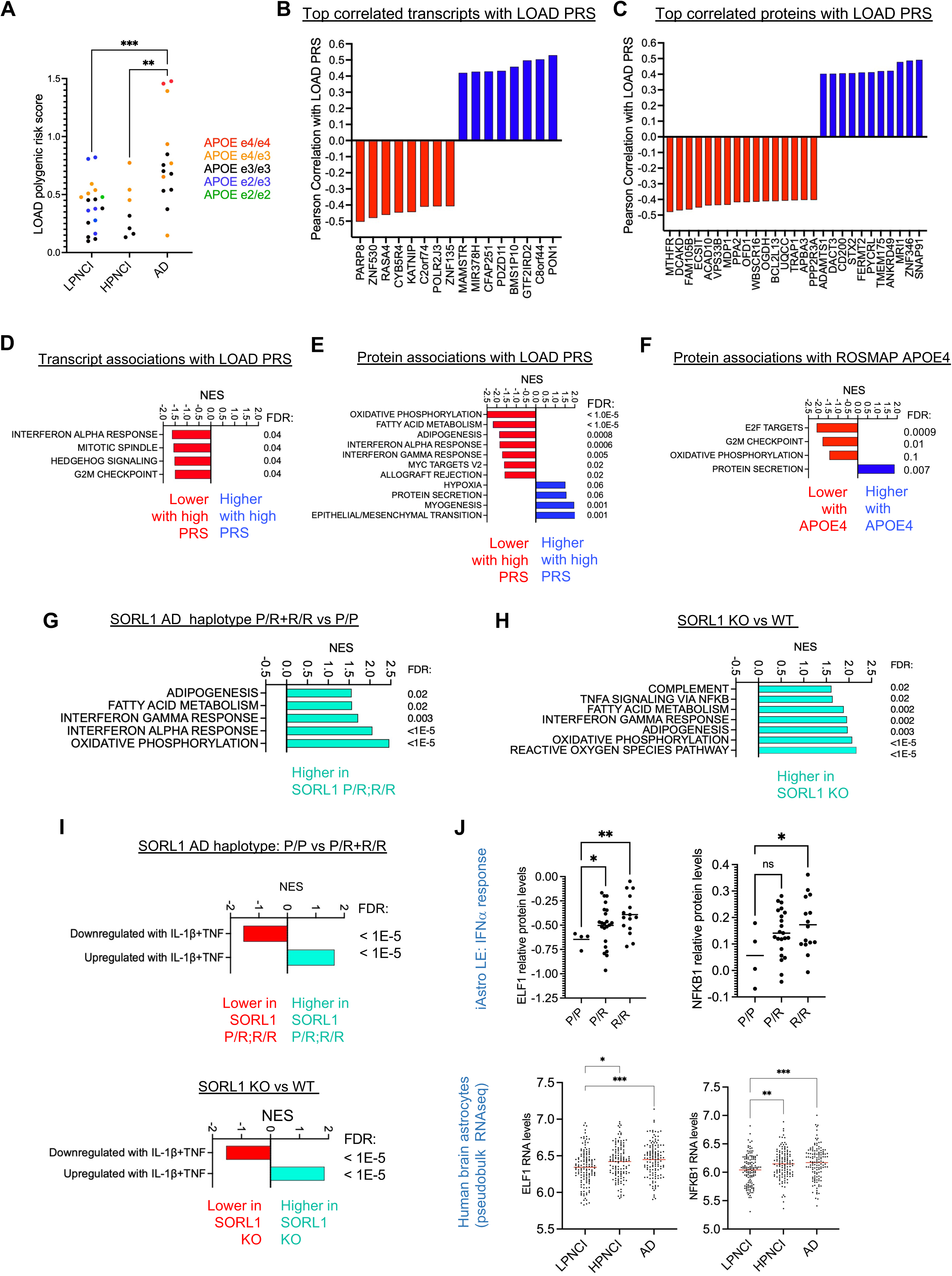
Genetic associations with RNA and protein profiles in iAs. (A) LOAD PRSes were calculated from genome sequencing data from individuals in ROSMAP using GWAS summary statistics (Bellenguez et al., 2022). Shown are the PRS distribution across categories for the iA cohort studied here, colored by APOE genotype. (B,C) Pearson correlation coefficients were calculated between LOAD PRS and each transcript quantified via RNAseq and each protein quantified by TMT-MS across iAs; shown are waterfalls of top associations. (D-G) GSEA was performed using all quantified genes (D) and proteins (E) to determine Hallmark pathways associated with LOAD PRS. (F) Proteomic profiles of ROSMAP iAs from e4 carriers were compared to iAs without e4, and GSEA performed to determine pathways enriched in iAs with e4. (G) ROSMAP iA proteomics data was analyzed to compare profiles of iAs with a SORL1 risk haplotype (P/R or R/R) and iAs with a homozygous protective haplotype (P/P). GSEA was performed using the Hallmark gene set to identify pathways associated with SORL1 risk haplotype. In parallel, isogenic SORL1 KO and WT iA protein profiles were compared, and GSEA performed using the same gene set. Shown in (G,H) are the concordant pathways enriched in both risk haplotype and SORL1 KO iAs. GSEA also was performed using the custom gene sets generated herein for genes up and down regulated in iAs with TNF + IL-1β stimulation (I). Two example LEPs upregulated with SORL1 risk alleles are shown that also are upregulated in AD brain tissue (J). For all panels, *p<0.05, **p<0.01, ***p<0.005.

After *APOE, SORL1* is one of the strongest genetic risk factors for AD (Alzheimer’s Association, 2023). SNPs at the *SORL1* locus have been repeatedly associated with LOAD in all large GWAS datasets, and rare autosomal dominant coding mutations in *SORL1* that affect its function have been identified in several families (Bellenguez et al., 2022; Kunkle et al., 2019; Lambert et al., 2013; Rogaeva et al., 2007; Vardarajan et al., 2015). We recently examined the consequences of loss of SORL1 in iAs, iNs, iMGs and iECs, and found that the loss of SORL1 had the greatest consequences on astrocyte and neuron RNA and protein profiles (Lee et al., 2023). Here, we examined associations of *SORL1* LOAD risk haplotype (Rogaeva et al., 2007) on ROSMAP iA profiles. With iAs, the presence of one (P/R) or two (R/R) copies of the risk haplotype was associated with higher levels of proteins involved in oxidative phosphorylation, fatty acid metabolism and interferon responses (**Fig. 8G**). In accord with an elevation in inflammation-related processes, genes found to be upregulated with TNF + IL-1β stimulation were upregulated in risk haplotype carriers, and those genes downregulated with stimulation were reduced in risk haplotype carriers (**Fig. 8I, Supp Tables 27,28**). We then performed these same analyses in isogenic *SORL1* KO versus WT iAs and found a concordant increase in these same pathways (**Fig. 8H,I, Supp Table 29,30**). Leading edge proteins upregulated with *SORL1* risk haplotype included ELF1 and NFKB1, two transcription factors critically linked to interferon and pro-inflammatory signaling ((Seifert et al., 2019); reviewed in (Anilkumar and Wright-Jin, 2024)). These transcription factors are also upregulated in astrocytes in the brain of individuals with AD (**Fig. 8J**). Collectively, these analyses highlight the utility of this resource for probing specific genetic variants across a large cohort of genetically diverse iA lines.

## Discussion

Molecular profiling of human brain from large, deeply phenotyped cohorts such as ROS and MAP have defined RNA-, protein-, and pathway level associations with amyloid and tau pathology in the brain, cognitive decline and resilience and with diagnosis of Alzheimer’s dementia and disease ((Johnson et al., 2022; Mostafavi et al., 2018) and others). Similarly, large scale single cell RNAseq data of postmortem ROSMAP brain also has revealed cell fate-specific RNA-level associations with these same traits ((Fujita et al., 2024; Green et al., 2023; Habib et al., 2020; Mathys et al., 2023) and others). These studies are quite valuable as they raise several mechanistic hypotheses based on data from a large number of humans. However, postmortem studies alone cannot distinguish between associations that represent causal elements of disease risk and progression and those that are consequences of decades of neuropathological aggregation and neuronal degeneration. Human experimental systems can play a key role in validating hypotheses and identifying therapeutic targets when coupled to deep phenotypic data from the humans from whom the cells were derived. Here, we describe a data and cell resource for interrogating the consequences of known and undefined genetic risk and resilience factors in astrocytes.

A recurring finding across both *in vitro* astrocytes and human brain tissue was the association between AD diagnosis and mitochondrial dysfunction, alongside a related association with fatty acid metabolism. IAs from individuals with AD exhibited reduced levels of proteins involved in oxidative phosphorylation, the TCA cycle, and fatty acid oxidation, which were similarly observed in brain tissue (**Fig. 6D,F**). Correspondingly, genetic analyses revealed that higher LOAD PRS was also associated with reduced levels of proteins involved in these processes (**Fig. 8E**), partially driven by the APOE genotype (**Fig. 8F**). Genes and proteins involved in protein trafficking and ER-Golgi sorting were upregulated in AD astrocytes and AD brain (**Fig. 6D,E**), and these were also associated with LOAD PRS and APOE genotype (**Fig. 8E,F**). These findings are logically interconnected, as LOAD PRS, APOE genotype, and AD diagnosis are all interrelated. However, it is important to recognize that these associations are not one-to-one. Approximately 35% of AD risk is estimated to be non-genetic (Sierksma et al., 2020), and PRS are based on GWAS summary statistics that do not represent an ethnically diverse population and do not capture the contributions of gene-disrupting rare mutations and structural variants. Thus, LOAD PRS does not fully predict AD (**Fig. 8A**). When associations are observed between pathways and LOAD PRS, it provides evidence that these findings are genetically driven by known genetic risk factors for AD.

An intriguing finding is the increase in basal interferon signaling observed in astrocytes from individuals resilient to AD neuropathology (**Fig. 7A-D**). Proteins upregulated in these resilient astrocytes include well-established interferon response genes including ISG20, ISG15, IFIT1, and factors directly involved in the transcriptional response to interferon including STAT2 and TRIM21 (**Fig. 7D**). We found that these resilient astrocytes secreted elevated levels of IFNγ, even without a stimulus (**Fig. 7G**). This suggests the presence of genetically-encoded protective factors that mildly increase basal interferon signaling, perhaps preparing these astrocytes to respond to insults like high extracellular Aβ. In line with a protective effect of elevated interferon signaling, low LOAD PRS was associated with elevated proteins involved in interferon signaling. Intriguingly, a recent study found that *APOE4* genotype is associated with elevation in cholesterol esters and reduced interferon-dependent pathways in human astrocytes, further implicating higher AD genetic risk with reduced interferon signaling (Feringa et al. 2024). Interferon signaling has been implicated in AD in many studies, for example, an elevation in interferon signaling is observed across mouse models of Aβ pathology (Roy et al., 2020). Multiple lines of evidence implicate that the activation of cyclic GMP-AMP synthase (cGAS)-“stimulator of interferon genes” (STING), results in microglia activation in the AD brain and AD animal models (reviewed in (Huang et al., 2023)). A recent study characterized “interferon response reactive astrocytes” using transcriptomic and proteomic profiling and identified genes and proteins specific to these astrocytes that also are enriched in AD mouse models and human AD postmortem brains (Hasel et al. 2021, Prakash et al. 2024). Together, these findings highlight the importance of further studies to identify what genetic resilience factors drive the elevation in interferon signaling in ROSMAP HP-NCI astrocytes, and to define the specific components of interferon signaling that are important for astrocyte function and astrocyte-microglia-neuron crosstalk.

*SORL1* has recently emerged as a fourth “causal” gene in AD pathogenesis. As noted above, GWAS have consistently identified LOAD-associated SNPs at the *SORL1* locus, and several coding mutations linked to early-onset AD have been found to reduce SORL1 levels. We recently showed that astrocytes, neurons, microglia, and endothelial cells each are affected by loss of SORL1 in both shared and divergent manners (Lee et al., 2023). Here, we show that the presence of the *SORL1* risk haplotype in iAs is associated with an elevation in proteins involved in oxidative phosphorylation, fatty acid metabolism and interferon signaling, findings mirrored in SORL1 KO astrocytes (**Fig. 8G,H**). Intriguingly, both *SORL1* risk haplotype and KO phenocopy the protein profile of astrocytes treated with IL-1β and TNF (**Fig. 8I,J**). These findings suggest that SORL1 plays a role in modulating basal signaling relevant to inflammatory processes in astrocytes.

A limitation of this study is that while it provides a molecular basis for defining the contribution of genetic variation in astrocytes to cell and molecular mechanisms, validation of these associations in an independent cohort of iPSC lines would further strengthen confidence in the findings and would provide enhanced power to define genetic associations for weaker genetic variants. Further, here we demonstrate how this highly reductionist system of pure astrocytes can be useful when coupled to data from the affected organ (in this case the brain). However, we anticipate that similar analyses in more complex human cellular models such as co-culture of neurons, astrocytes, microglia and vascular cells from this iPSC cohort will be highly valuable for building on this work to capture and uncover a large proportion of the contribution of genetic variation to AD. Further, the addition of AD- and aging-relevant stressors (such as Aβ and human AD brain extracts, (Hsieh et al., 2022; Jin et al., 2018)) also will enhance the experimental model to further interrogate the potential for astrocytes to contribute to risk and resilience. Single cell analyses revealed three subtypes of iAs that each responded to pro-inflammatory factors in overlapping but distinct manners (**Fig. 4**). Co-culture with other cell types as well as introduction of stressors into the model system may further enable the modeling of additional astrocyte subtypes found in the postmortem human brain (Fujita et al., 2024; Green et al., 2023; Habib et al., 2020; Mathys et al., 2023). This iPSC collection and the associated datasets developed here for astrocytes and previously described for neurons (Lagomarsino et al., 2021) can serve as a resource for future studies aimed at disentangling the contribution of genetic variation to cellular processes affecting risk and resilience to LOAD.

## Acknowledgements

We thank the NeuroTechnology Studio at Brigham and Women’s Hospital for providing LSM710 and Chromium 10x instrument access and consultation on data acquisition and data analysis; D. Duong (Emory Integrated Proteomics Core at Emory University School) for technical assistance with TMT-MS; F. Quintana, T. Schwartz, and D. Selkoe for their Dissertation Advisory Committee guidance; members of the Young-Pearse lab for their critical reading and manuscript feedback. This work was supported by NIH grants F31AG063399, U01AG46152, U01AG61356, U01AG072572, RF1AG057473, and R01AG063398. The results published here are in part based on data obtained from the AD Knowledge Portal (https://adknowledgeportal.org). ROS and MAP study data were provided by the Rush Alzheimer’s Disease Center, Rush University Medical Center, Chicago. Data collection was supported through funding by NIA grants P30AG10161 (ROS), R01AG15819 (ROSMAP; genomics and RNAseq), R01AG17917 (MAP), R01AG36836 (RNAseq), RF1AG57473 (single nucleus RNAseq), U01AG32984 (genomic and whole exome sequencing), the Illinois Department of Public Health (ROSMAP), and the Translational Genomics Research Institute (genomic). Additional phenotypic data and iPSC lines can be requested at www.radc.rush. edu. For TMT Proteomics of brain tissue, study data were provided through the Accelerating Medicine Partnership for AD (U01AG046161 and U01AG061357) based on samples provided by the Rush Alzheimer’s Disease Center, Rush University Medical Center, Chicago. Data collection was supported through funding by NIA grants R01AG30146, R01AG36836, U01AG46152, the Illinois Department of Public Health, and the Translational Genomics Research Institute. For snRNAseq data, study data were generated from postmortem brain tissue provided by the ROSMAP cohort at Rush Alzheimer’s Disease Center, Rush University Medical Center, Chicago. This work was funded by NIH grants U01AG061356, RF1AG057473 and U01AG046152 as part of the AMP-AD consortium, as well as NIH grants R01AG066831 and U01AG072572.

---

**Supp. Fig. 1.**
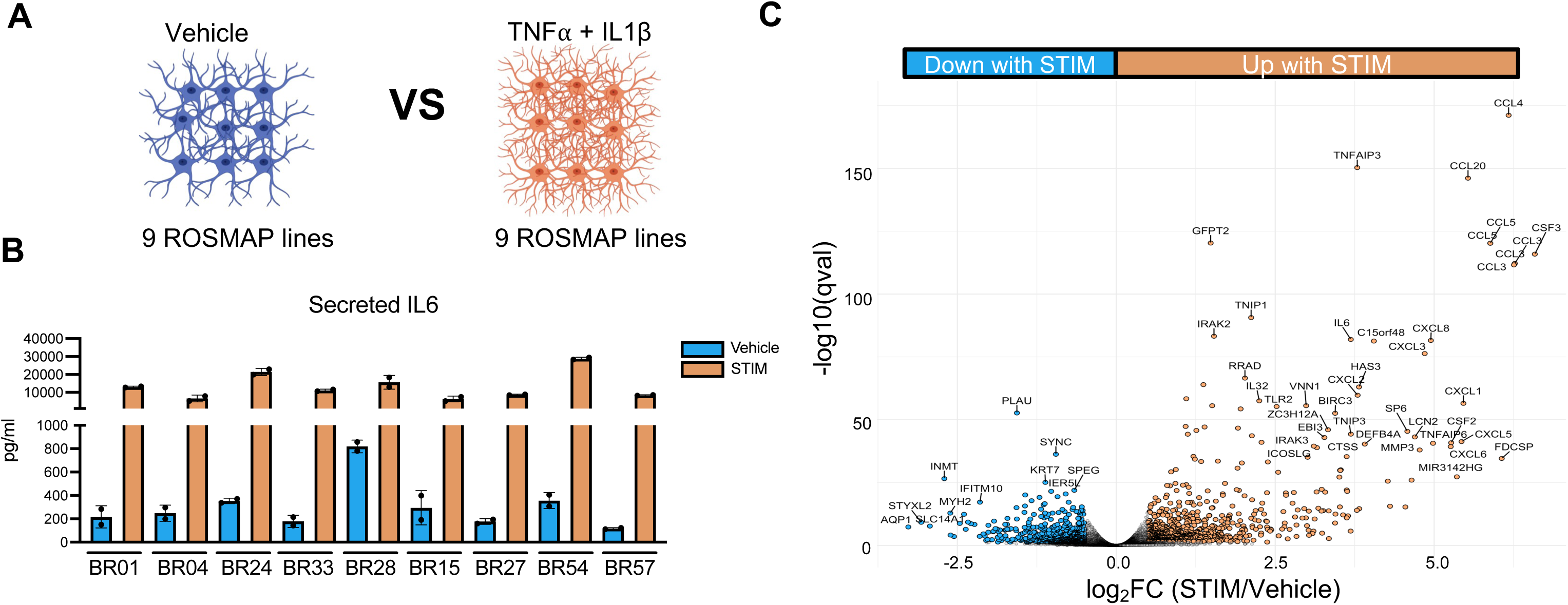
Characterizing iA response to cytokine stimulation. (A) Schematic of the experimental design. 24hrs prior to harvest, iAs from the 9 lines were treated with either vehicle (0.1% BSA in PBS) or cytokines (5ng/ml TNF and 1ng/ml IL-1β) and conditioned media and RNA samples were collected. (B) Secreted levels of proinflammatory cytokine IL-6 were measured using MSD Proinflammatory cytokines ELISA. Each dot represents a value from unique cell line, values are normalized to protein concentration. Data show mean +/− SEM. (C) Volcano plot of genes differentially expressed with stimulation Adjusted p value (q value) <0.05 and absolute logFC value >0.5 were used as cutoff values.

**Supp. Fig. 2.**
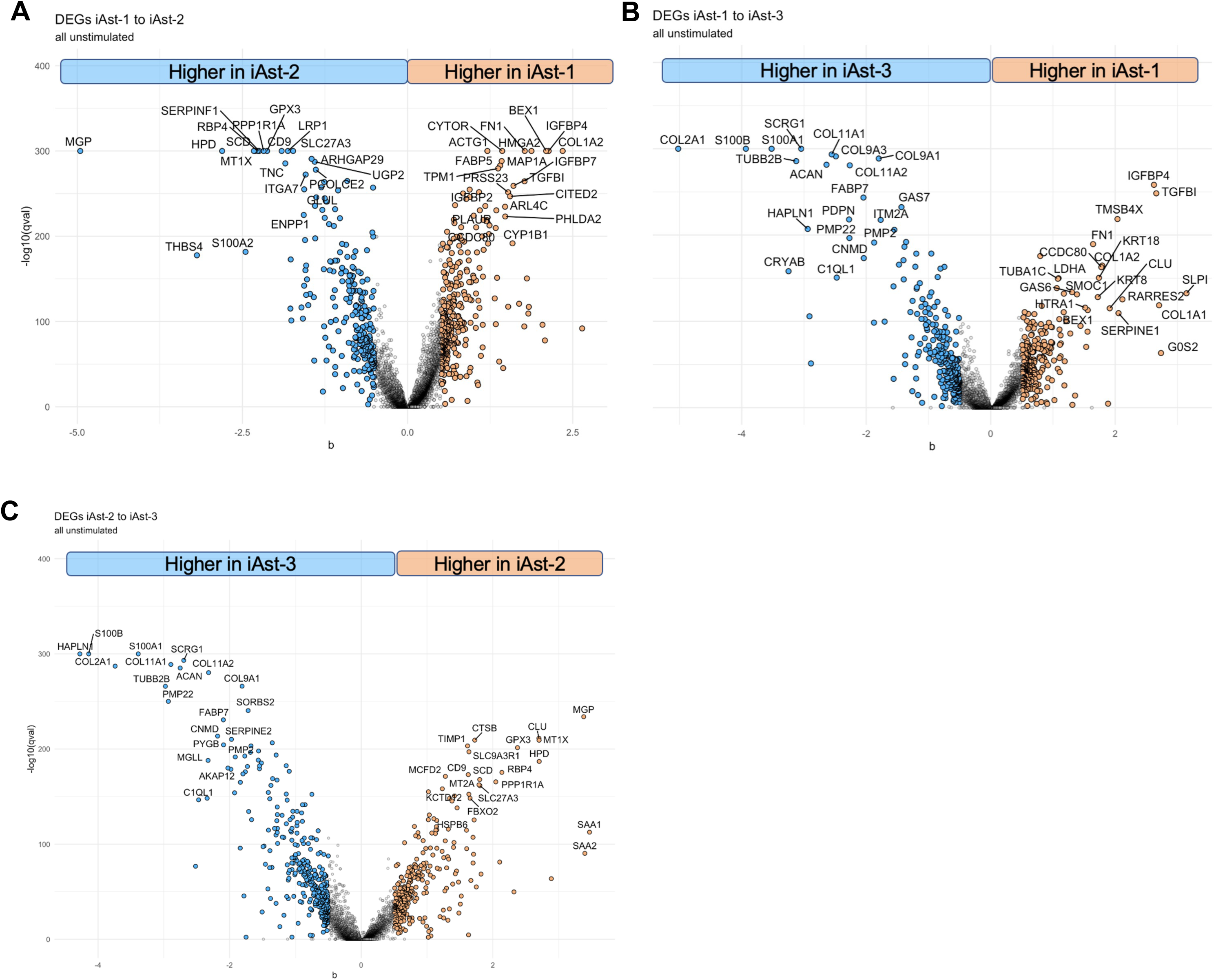
Differentially Expressed Genes across different iA clusters. (A-C) Volcano plot of genes differentially expressed between (A) iAst 1 vs 2 (B) iAst 1 vs 3 and (C) iAst 2 vs 3.

**Supp. Fig. 3.**
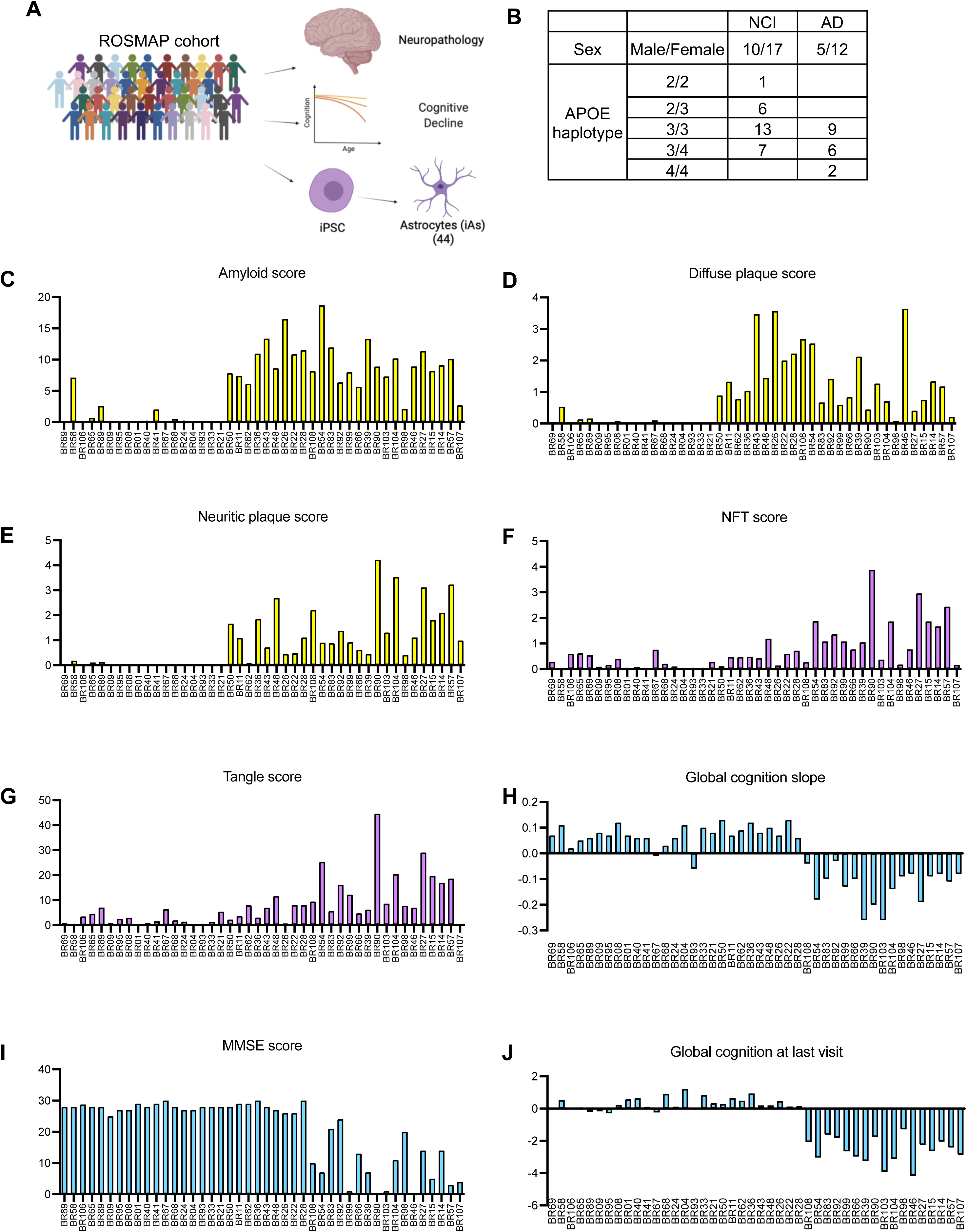
ROSMAP cohort characterization. (A) Schematic of the experimental design. Based on neuropathological and clinical data, 44 individuals were selected. Then, PBMCs were collected from ROSMAP cohort participants, which were converted to iPSCs and differentiated into astrocytes (iAs). NCI = not cognitively impaired, AD = Alzheimer’s disease. (B) Table showing sex and APOE haplotypes of individuals in each category. (C-J) Summary of global pathology score, global cognition at last visit, and age at death for 44 individuals used in this study. One-way ANOVA with Tukey’s multiple comparisons test. (C) Amyloid score (D) Diffuse plaque score (E) Neuritic plaque score (F) Neurofibrillary tangles (NFT) score (G) Tangle score (H) Global cognition slope. (I) Mini-mental state examination (MMSE) score (J) Global cognition at last visit (C-J) These variables have been previously defined in (Lagomarsino et al., 2021).

**Supp. Fig. 4.**
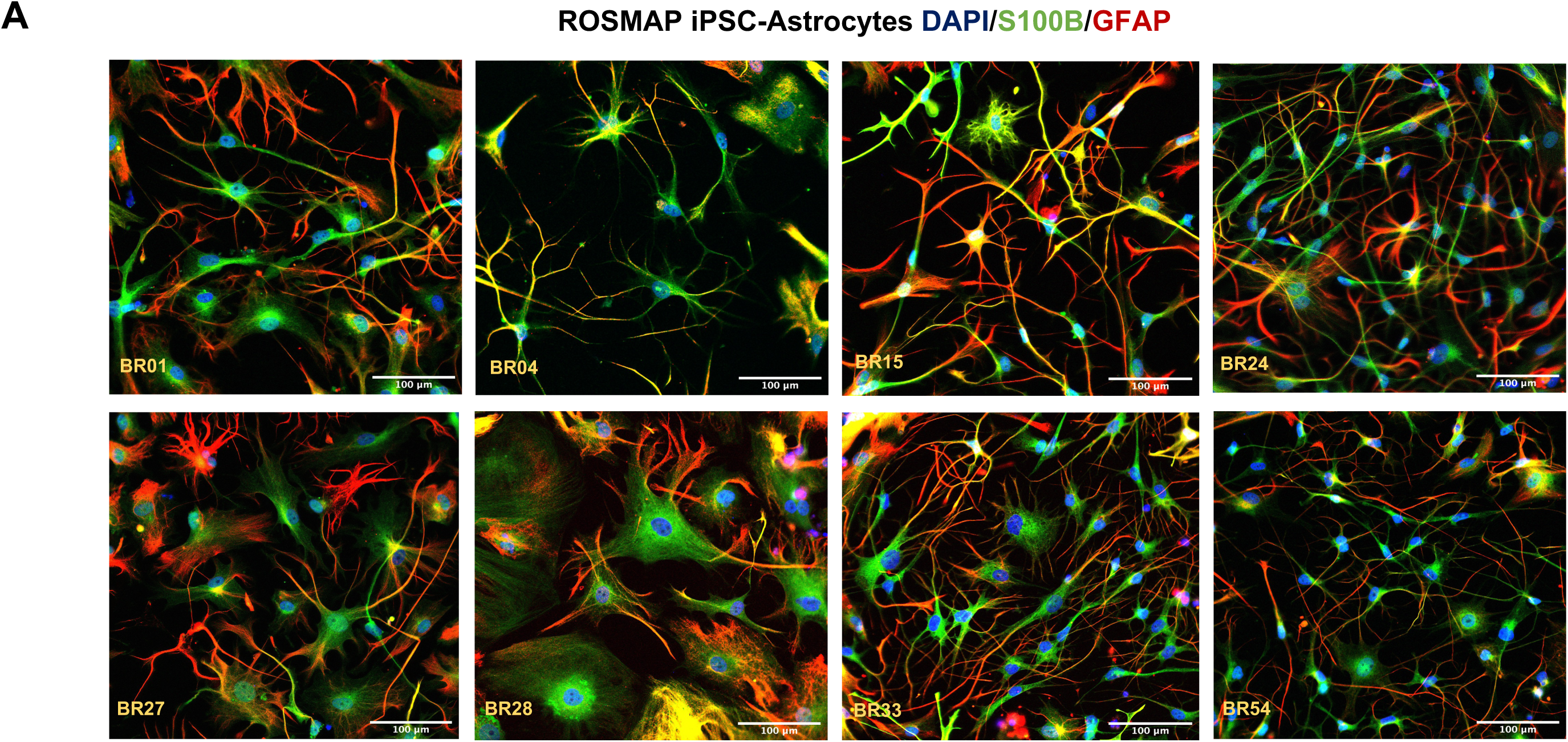
Representative immunostaining of iAs. iAs generated from ROSMAP iPSCs were immunostained for GFAP and S100B. Representative images for eight lines are shown. Scale bar = 100 μm.

**Supp. Fig. 5:**
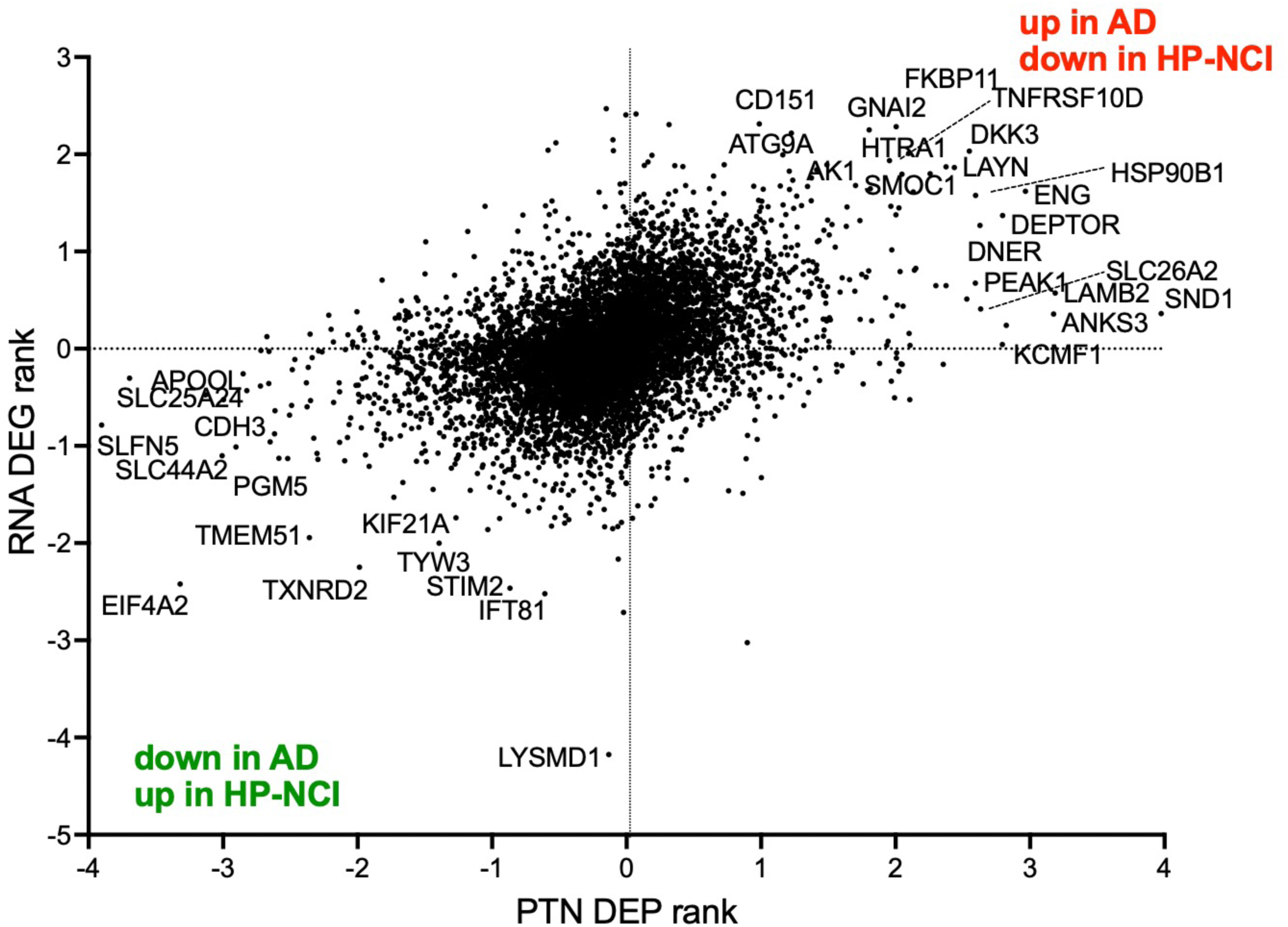
Integration plot of genes and proteins differentially expressed in resilient vs AD iAs. (A) Results of proteomics comparison of HPNCI vs AD iAs are plotted along the x-axis and results of RNAseq comparison of HPNCI vs AD iAs are plotted along the y-axis to determine concordant changes at the gene and protein level. Values are the ranks (−log10 (p-value); signed based upon direction of change) following DEP and DEG analyses.

**Supp. Fig. 6.**
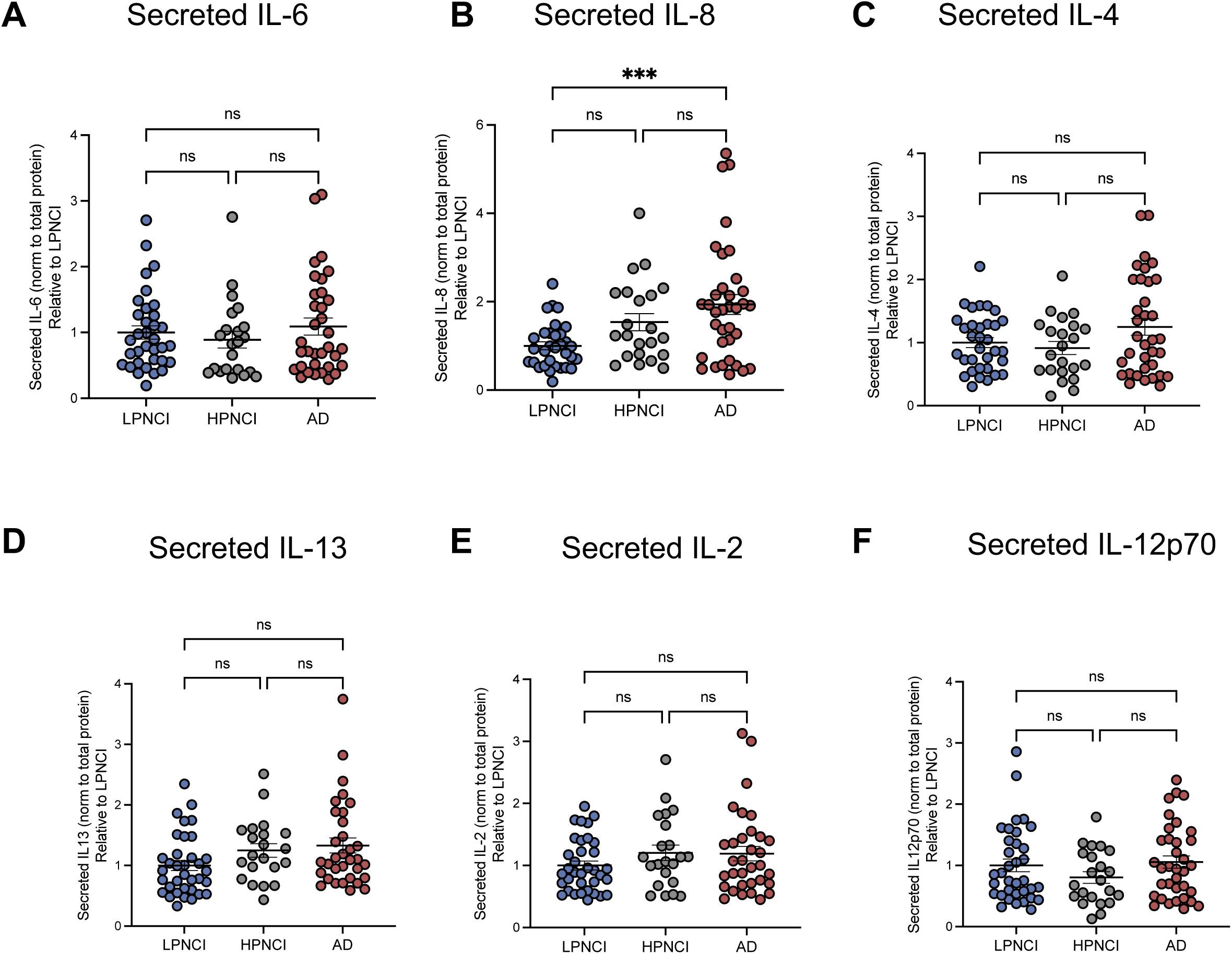
Characterization of proinflammatory cytokines secreted from iAs. (A) Additional cytokines analysis from Figure 8B. Secreted levels of IL-6, IL-8, IL-4, IL-13, IL-2, and IL-12p70 were measured via ELISA and categorized by diagnosis. Each measurement is normalized to total protein and values shown are relative to LPNCI individuals. One-way ANOVA with Tukey’s multiple comparison’s test, n=2 per cell line.

